# Structures of the Catalytically Activated Yeast Spliceosome Reveal the Mechanism of Branching

**DOI:** 10.1101/500363

**Authors:** Ruixue Wan, Rui Bai, Chuangye Yan, Jianlin Lei, Yigong Shi

## Abstract

Pre-mRNA splicing is executed by the spliceosome. Structural characterization of the catalytically activated complex (B^*^) is pivotal for mechanistic understanding of catalysis of the branching reaction by the spliceosome. In this study, we assembled the B^*^ complex on two different pre-mRNAs from *Saccharomyces cerevisiae* and determined the cryo-EM structures of four distinct B complexes at overall resolutions of 2.9-3.8 Å. The duplex between U2 snRNA and the branch point sequence (BPS) is located 13-20 Å away from the 5’-splice site (5’SS) in the B^*^ complexes that are devoid of the step I splicing factors Yju2 and Cwc25. Recruitment of Yju2 into the active site brings the U2/BPS duplex into the vicinity of 5’SS, ready for branching. In the absence of Cwc25, the nucleophile from BPS is positioned about 4 Å away from, and remains to be activated by, the catalytic metal M2. This analysis reveals the functional mechanism of Yju2 and Cwc25 in branching. These four structures constitute compelling evidence for substrate-specific conformations of the spliceosome in a major functional state.

## Introduction

Pre-mRNA splicing is executed by the spliceosome (Brody and Abelson, 1985; Frendewey and Keller, 1985; Grabowski et al., 1985), a supramolecular complex with exceptional dynamics in its composition and conformation (Shi, 2017a; Will and Luhrmann, 2011). The fully assembled spliceosome exists in at least eight major functional states: precursor of the pre-catalytic spliceosome (pre-B), pre-catalytic spliceosome (B), activated complex (B^act^), catalytically activated complex (B^*^), catalytic step I complex (C), step II activated complex (C^*^), post-catalytic complex (P), and intron lariat spliceosome (ILS). Each splicing cycle, involving branching and exon ligation, results in the removal of the intervening RNA sequences between two target exons (Grabowski et al., 1984; Padgett et al., 1984; Ruskin et al., 1984). Branching and exon ligation proceed spontaneously in the B^*^ and C^*^ complexes, respectively (Shi, 2017b). The B^act^-to-B^*^ and C-to-C* transitions, however, are driven by the ATPase/helicases Prp2 and Prp16, respectively (Cordin et al., 2012; Jankowsky, 2011; Liu and Cheng, 2015).

The first near-atomic resolution structure of an assembled spliceosome, determined at 3.6 Å by single-particle cryo-EM analysis, was reported for the ILS complex from *S.pombe* in 2015 (Hang et al., 2015; Yan et al., 2015). Since then, 13 cryo-EM structures, mostly at resolutions between 3.3 and 5.8 Å, have been elucidated for seven distinct states of the assembled spliceosome from *S. cerevisiae* (Bai et al., 2018; Bai et al., 2017; Fica et al., 2017; Galej et al., 2016; Liu et al., 2017; Plaschka et al., 2017; Rauhut et al., 2016a; Wan et al., 2016b; Wan et al., 2017; Wilkinson et al., 2017; Yan et al., 2016, 2017). Seven such structures have been reported for five distinct states of the human spliceosome (Bertram et al., 2017a; Bertram et al., 2017b; Haselbach et al., 2018; Zhan et al., 2018a, b; Zhang et al., 2017; Zhang et al., 2018). Among the eight known functional states of the spliceosome, only the B^*^ complex remains structurally uncharacterized.

In this study, we report the cryo-EM structures of the B^*^ complex from *S. cerevisiae.* The B^*^ complex was prepared using two different pre-mRNAs, which give rise to an average resolution of 2.9 Å for the B^*^ complex assembled on the *ACT1* pre-mRNA and 3.2 Å for that assembled on the *UBC4* pre-mRNA. For each of the two distinct B^*^ complexes, two different conformational states were observed. Comparison of these conformational states reveals mechanistic insights into the branching reaction.

## Results

### Preparation and EM analysis of the B^*^ complex

It is difficult to isolate the highly transient B^*^ complex. Relying on direct purification of the spliceosomes from cells, we have been unable to purify the endogenous B^*^ complex. We sought to reconstitute the B^*^ complex using the *in vitro* assembly approach, in which the assembled B^act^ complex was remodeled into the B^*^ complex upon incubation with recombinant Prp2 and Spp2 in the presence of ATP (Bao et al., 2017; Ohrt et al., 2013; Roy et al., 1995; Warkocki et al., 2009; Warkocki et al., 2015) (Figure S1A-C). To maximize our chance of success, we used two different pre-mRNA: *ACT1* and *UBC4* (Figure S1C). The 3’-end nucleotides of the 5’-exon in *ACT1* are UCUG; in contrast, the corresponding nucleotides in *UBC4* are AAAG, which may make improved base-pairing interactions with the poly-U sequences of U5 loop I. In both cases, we were able to isolate the spliceosomes that appeared to be intact by negative staining EM analysis (Figure S1D,E).

For both samples, we prepared the cryo-EM grids and performed EM analysis (Figure S1F-H). 1.8 and 1.7 million particles were auto-picked for the spliceosomes assembled on the *ACT1* and *UBC4* pre-mRNA, respectively (Figures S2 & S3). Following two rounds of three-dimensional (3D) classification, 555,036 particles yielded a final reconstruction of the B^*^ complex at an average resolution of 2.9 Å (Figures S2 & S4; Tables S1 & S2). Despite excellent EM density for much of the *ACT1* B^*^ complex (Figures S5), the density in the 5’SS region is weak. Next, we processed the data on the spliceosomes assembled on *UBC4.* After two rounds of 3D classification, 132,125 particles yielded a final structure of the *UBC4* B^*^ complex at an average resolution of 3.2 Å (Figures S3, S4, S6; Tables S1 & S3). Use of *UBC4* indeed led to improved EM density in the 5’SS region.

For each of the two B^*^ complexes, the local resolution in the U2/BPS duplex region is relatively low. This is likely due to absence of the step I splicing factors Cwc25 and Yju2, which may stabilize the conformation of the nucleophile as well as the acceptor (Galej et al., 2016; Wan et al., 2016b). To improve the local density, we applied a soft mask on the U2/BPS region and performed additional 3D classification. This effort led to two distinct conformational states of the *ACT1* B^*^ complex at average resolutions of 3.6 and 3.2 Å, named B^*^-a1 and B^*^-a2, respectively (Figure 1A; Figures S2 & S7A,B). A similar analysis on the *UBC4* B^*^ complex also resulted in two distinct conformations at 3.9 and 3.7 Å resolution, designated as B^*^-b1 and B^*^-b2, respectively (Figure 1B; Figures S3 & S7C,D).

**Figure 1.**
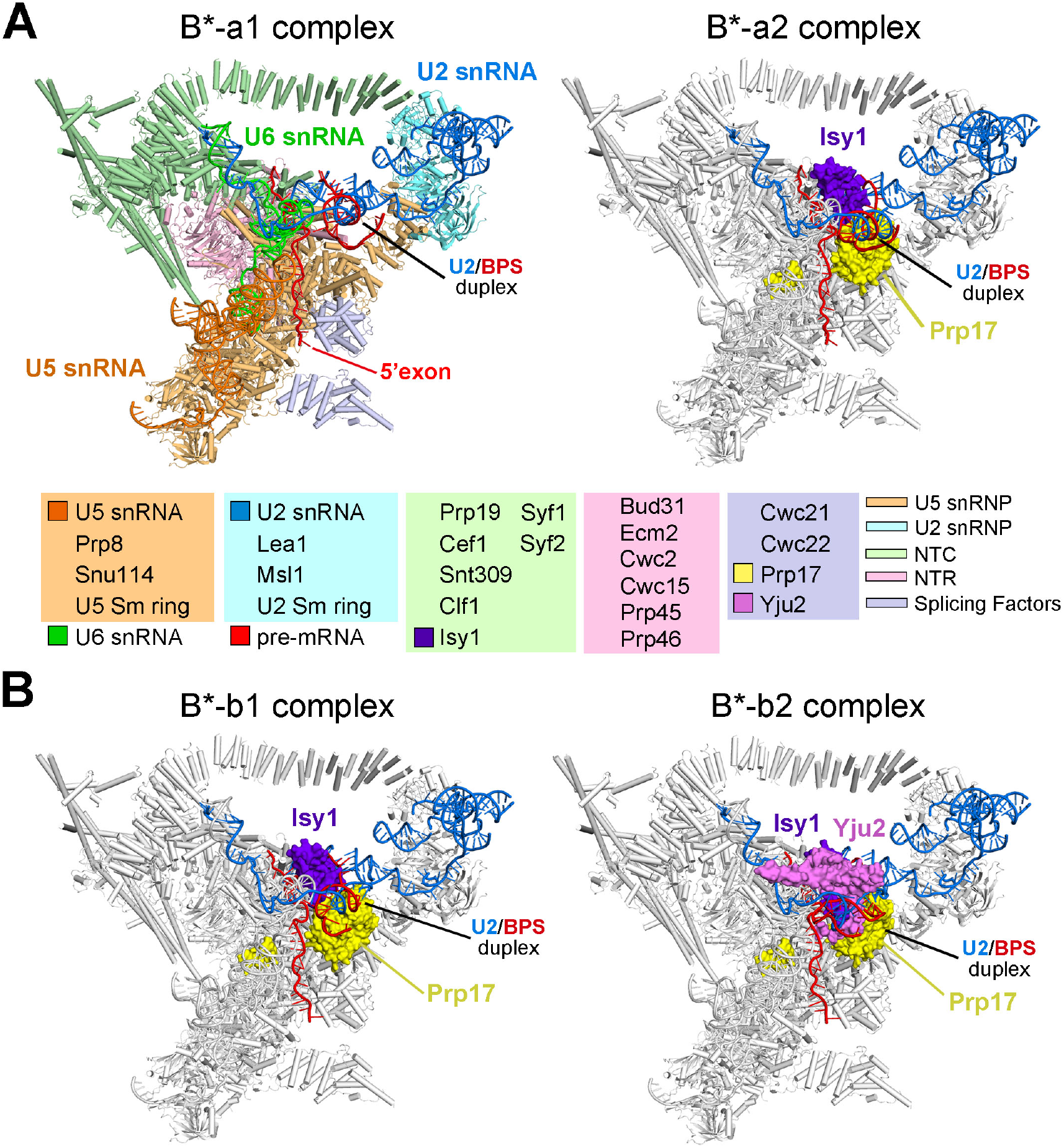
Cryo-EM structures of the catalytically activated spliceosomes (B^*^ complexes) from *Saccharomyces cerevisiae (S. cerevisiae).* (A) Structures of the B^*^ complex assembled on the *ACT1* pre-mRNA. Two distinct conformational states were captured and named B^*^-a1 (left panel) and B^*^-a2 (right panel). The final atomic model includes 34 proteins, three snRNAs, and 60 nucleotides of the *ACT1* pre-mRNA. The snRNPs and major components of the B^*^-a1 complex are color-coded. The pre-mRNA, U2, U5 and U6 snRNAs are colored red, marine, orange and green, respectively. The splicing factors are colored light purple. Compared to B^*^-a1, only those components that undergo changes are colored in the B^*^-a2 complex, with the rest shown in grey. The protein and RNA components are tabulated below the structures. (B) Structures of the B^*^ complex assembled on the *UBC4* pre-mRNA. Two distinct conformational states were captured and named B^*^-b1 (left panel) and B^*^-b2 (right panel). Compared to B^*^-a1, only those components that undergo changes are colored in these complexes. Compared to the other three B^*^ complexes, B^*^-b2 contains an extra protein – the step I splicing factor Yju2 (violet). All structural images were created using PyMol (DeLano, 2002).

### Structure of the B^*^ complex

Except for Yju2 in B^*^-b2, all four B^*^ complexes share the same set of RNA and protein components (Figure 1). The overall structures of these components are very similar among the four B^*^ complexes except for the step II splicing factor Prp17, the NTC component Isy1, and the local conformation of two RNA elements: the U2/BPS duplex and the 5’-exon-5’SS sequence. The final atomic model for each of the four B^*^ complexes from *S. cerevisiae* contains a pre-mRNA, U6 small nuclear RNA (snRNA), U5 small nuclear ribonucleoprotein (snRNP), NineTeen complex (NTC), NTC-related (NTR), U2 snRNP core (U2 snRNA, Leal, Msl1, and the U2 Sm ring), and the splicing factors Cwc21, Cwc22 and Prp17 (Figure 1A). The shared components of these four B^*^ complexes include 34 proteins and four RNAs (Tables S1-S3). The WD40 domain of Prp17 is invisible only in the B^*^-a1 complex but is clearly identified in the other three B^*^ complexes; it is positioned in slightly different locations in these complexes, presumably due to the relative movements of the U2/BPS duplex.

Replacement of *ACT1* by *UBC4* in the B^*^ complex led to improvement of the EM density for the RNA elements in the splicing active site center (Figure S6B,C). Consequently, base-pairing interactions among the four pieces of RNA are unambiguously assigned in the *UBC4* B^*^ complex (Figure S7E). The step I splicing factor Yju2, which is thought to be recruited into the B^act^ complex along with the NTC components (Chang et al., 2009; Liu et al., 2007; Warkocki et al., 2009), was not structurally identified in the B^act^ complex (Rauhut et al., 2016b; Yan et al., 2016). Notably, however, Yju2 is present only in the B^*^-b2 complex, but not in the other three B^*^ complexes (Figure 1B). The N-terminus of Isy1 is positioned only in the active site of the B^*^-b2 complex but disordered in the other three complexes, likely due to the presence of Yju2 in B^*^-b2. A total of 548 nucleotides have been identified in the *UBC4* B^*^ complex (Figure 2A). Among these, 491 are assigned to the three snRNAs and 57 to the *UBC4* pre-mRNA.

### The RNA elements in the B^*^ complex

A major difference among the four B^*^ complexes is the conformation and the location of the key RNA elements in the catalytic center (Figures 1 & 2). These RNA elements include the U2/BPS, 5’SS, and U5/5’-exon duplexes. The conformation and precise positioning of the U2/BPS duplex in the active site require stabilization by the step I factors Cwc25 and Yju2 (Galej et al., 2016; Schneider et al., 2015; Tseng et al., 2017; Wan et al., 2016b). In the absence of Cwc25 and Yju2, the U2/BPS duplex is not positioned in the close vicinity of the 5’SS and its exact conformation and location differ among the three B^*^ complexes: a1, a2, and b1 (Figure 2B,C). The nucleophile from the BPS is located about 20 Å away from the phosphate of G_1_ in the 5’SS in the B^*^-a1 complex (Figure 2B, left panel); this distance is shortened to approximately 13 Å in B^*^-a2 (Figure 2B, right panel) and 15 Å in B^*^-b1 (Figure 2C, left panel). In sharp contrast, the U2/BPS duplex is translocated to the vicinity of the 5’SS in the Yju2-containing B^*^-b2 complex, with the 2’-OH of BPS A_70_ poised for nucleophilic attack on the phosphate of G_1_ in the 5’SS that is only 4.3 Å away (Figure 2C, right panel).

**Figure 2.**
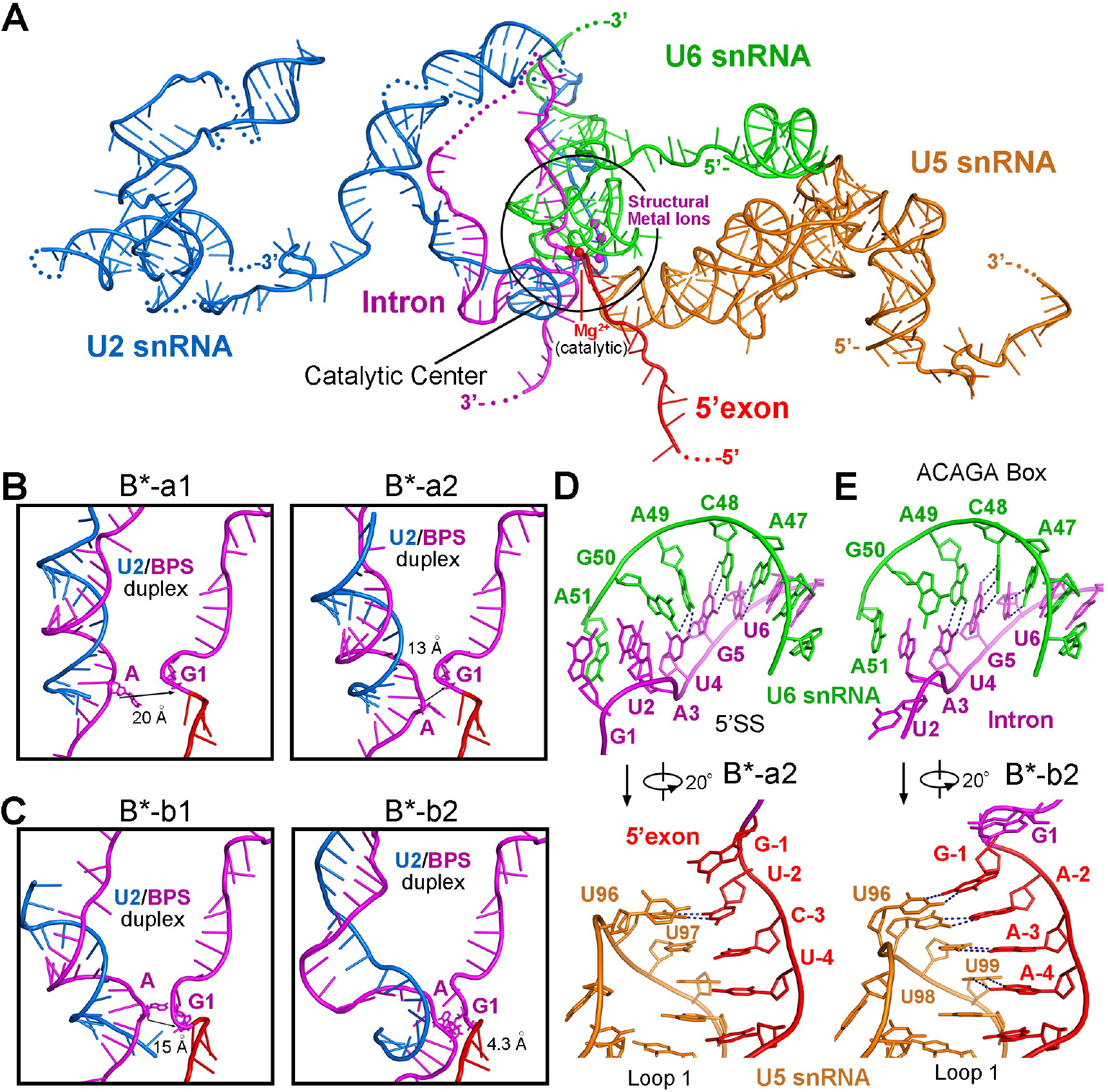
The RNA elements in the *S. cerevisiae* B^*^ complexes. (A) Structure of the RNA elements in the B^*^-b2 complex. The *UBC4* pre-mRNA and U2, U5, and U6 snRNAs are colored red, marine, orange, and green, respectively. The catalytic and structural metal ions are shown as red and magenta spheres, respectively. (B) Comparison of the U2/BPS duplexes from the B^*^-a1 and B^*^-a2 complexes. The nucleophile is located about 20 and 13 Å away from the acceptor in the B^*^-a1 and B^*^-a2 complexes, respectively. (C) Comparison of the U2/BPS duplexes from the B^*^-b1 and B^*^-b2 complexes. In the B^*^-b1 complex (left panel), the nucleophile is about 15 Å away from the acceptor. In the B^*^-b2 complex (right panel), the nucleophile is positioned 4 A away and almost ready for the branching reaction. (D) A close-up view on the 5’SS/U6 duplex (upper panel) and the 5’-exon/U5 duplex (bottom panel) in the B^*^-a2 complex. The 5’SS is recognized by the ACAGA box of U6 snRNA through base-paring interactions between U_4_G_5_U_6_ of pre-mRNA and A_47_C_48_A_49_ of U6 snRNA. The 5’-exon is recognized by loop I of U5 snRNA, with U_-2_ of 5’-exon base-pairing with U_97_ of U5 snRNA. (E) A close-up view on the structure of the 5’SS/U6 duplex (upper panel) and the 5’-exon/U5 duplex (bottom panel) in the B^*^-b2 complex. The 5’SS is recognized identically as that in the B^*^-a2 complex; the 5’-exon, however, is now anchored to loop I of U5 snRNA through base-pairing interactions between A_-4_A_-3_A_-2_G_-1_ of the 5’-exon and U_96_U_97_U_98_U_99_ of U5 snRNA.

Recognition of the highly conserved 5’SS by U6 snRNA is nearly identical among the four B^*^ complexes. Specifically, three contiguous nucleotides U_4_G_5_U_6_ of the 5’SS form a duplex with A_47_C_48_A_49_ of the ACAGA box of U6 snRNA through canonical base-pairing interactions (Figure 2D,E, upper panels). On the other hand, different sequences of *ACT1* and *UBC4* at the 3’-end of their 5’-exons lead to recognition of different strengths by U5 snRNA. The nucleotides G_-1_U_-2_C_-3_U_-4_ of the *ACT1* 5’-exon are bound loosely to the loop I of U5 snRNA, with U_-2_ of *ACT1* base-pairing to U_96_ of U5 snRNA (Figure 2D, lower panel). In contrast, the nucleotides G_-1_A_-2_A_-3_A_-4_ of the *UBC4* 5’-exon form a duplex with U_96_U_97_U_98_U_99_ of U5 snRNA through canonical base-pairing H-bonds (Figure 2E, lower panel).

### The splicing active site and metal ions

In the B^*^-b2 complex, the splicing active site comprises loop I of U5 snRNA, helix I of the U2/U6 duplex, intramolecular stem loop (ISL) of U6 snRNA and five associated metal ions (Figure 3A). The nucleophile and the acceptor of the branching reaction are already delivered into the active site. Four consecutive nucleotides at the 3’-end of the 5’-exon are anchored to U5 loop I. Helix Ib and three nucleotides of U6 snRNA form a characteristic catalytic triplex (Figures S6B & S7E). Remarkably, the nucleophile-containing A_70_ of the BPS interacts with G_1_U_2_ of the 5’SS, with the adenine base making canonical H-bonds to the uracil while stacking against the guanine (Figure 3B). The nucleophile 2’-OH of A_70_ is located about 4.3 Å away from the phosphorous atom of G_1_, ready to be activated by a catalytic magnesium ion (M2) for the branching reaction.

Among the five metal ions, three are coordinated exclusively by the ISL and may play a structural role by neutralizing the negative charges of the RNA (Figure 3A). These three metals are identically positioned in all four B^*^ complexes. The other two metals, presumably Mg^2+^, have been identified as M1, which stabilizes the leaving group during the branching reaction, and M2, which activates the nucleophile (Fica et al., 2014; Fica et al., 2013; Keating et al., 2010; Steitz and Steitz, 1993; Yean et al., 2000). In the B^*^-a1 and -a2 complexes, M1 is coordinated by the phosphate oxygens of G_78_ and U_80_ from U6 snRNA and M2 is bound by a phosphate oxygen from U_80_ of U6 snRNA (Figure 3C,D). M1 is identically coordinated in the B^*^-b1 complex (Figure 3E) but is already bound to the phosphate oxygen of G_1_ of the 5’SS in the B^*^-b2 complex (Figure 3F). Compared to B^*^-a1/a2, M2 gains an additional ligand (a phosphate oxygen from G_60_ of U6 snRNA) in the B^*^-b1 complex (Figure 3E) and two additional ligands (two phosphate oxygens from A_59_ and G_60_ of U6 snRNA) in the B^*^-b2 complex (Figure 3F). But the 2’-OH of A_70_ is yet to be activated by M2, with a distance of approximately 4.3 Å. The observation that the phosphate of U_80_ of U6 snRNA coordinates both M1 and M2 in all four B^*^ complexes is consistent with the finding that sulfur substitution of one of the non-bridging phosphoryl oxygens in U_80_ abolishes the branching reaction (Fica et al., 2014; Yean et al., 2000).

Among the four B^*^ complexes, B^*^-b2 appears to be nearly ready for the branching reaction. In response to Yju2 binding, the 5’-exon-5’SS junction undergoes a major reorganization in the B^*^-b2 complex compared to the other three (Figure 3C-F). In particular, the phosphate group of G_1_ of the 5’SS is flipped inside out to coordinate the M1 metal. Consistent with published observations (Bao et al., 2017), such a dynamic change is afforded by the dissociation of Cwc24 and Prp11, which protect Gļ in the B^act^ complex (Rauhut et al., 2016b; Yan et al., 2016), and the recruitment of Yju2, which stabilizes the local conformation in the B^*^-b2 complex. The recruitment of Cwc25 may allow slight rearrangement of the active site RNA elements and consequent activation of the nucleophile by M2 (Chiu et al., 2009; Galej et al., 2016; Wan et al., 2016b; Warkocki et al., 2009).

### Structural changes of the B^act^-to-B^*^ transition

The B^act^-to-B^*^ transition requires the ATPase/helicase Prp2 and involves a major reorganization of the spliceosome (Bao et al., 2017; Kim and Lin, 1996; Lardelli et al., 2010; Ohrt et al., 2012; Warkocki et al., 2009; Warkocki et al., 2015) (Figure 4A). During this transition, at least 16 proteins are dissociated and several proteins undergo conformational changes. In the B^act^ complex, the SF3b complex and the RES complex recognize the U2/BPS duplex and the downstream RNA sequences of the intron, respectively; Prp11 from the SF3a complex and the splicing factor Cwc24 are placed in between the U2/BPS duplex and the 5’SS (Yan et al., 2016) (Figure 4A). Upon the action of Prp2, SF3a, SF3b, RES and the splicing factors Cwc24 and Cwc27 are dissociated to release the U2/BPS duplex and the 5’SS (Lardelli et al., 2010; Ohrt et al., 2012; Warkocki et al., 2009). U5 snRNP, NTR, and the splicing factors Cwc21 and Cwc22 remain static. In contrast, the U2 snRNP core and the superhelical proteins Syf1 and Clf1 of the NTC complex undergo marked translocation. These changes are similar to those described for the B^act^-to-C transition (Wan et al., 2016b).

**Figure 3.**
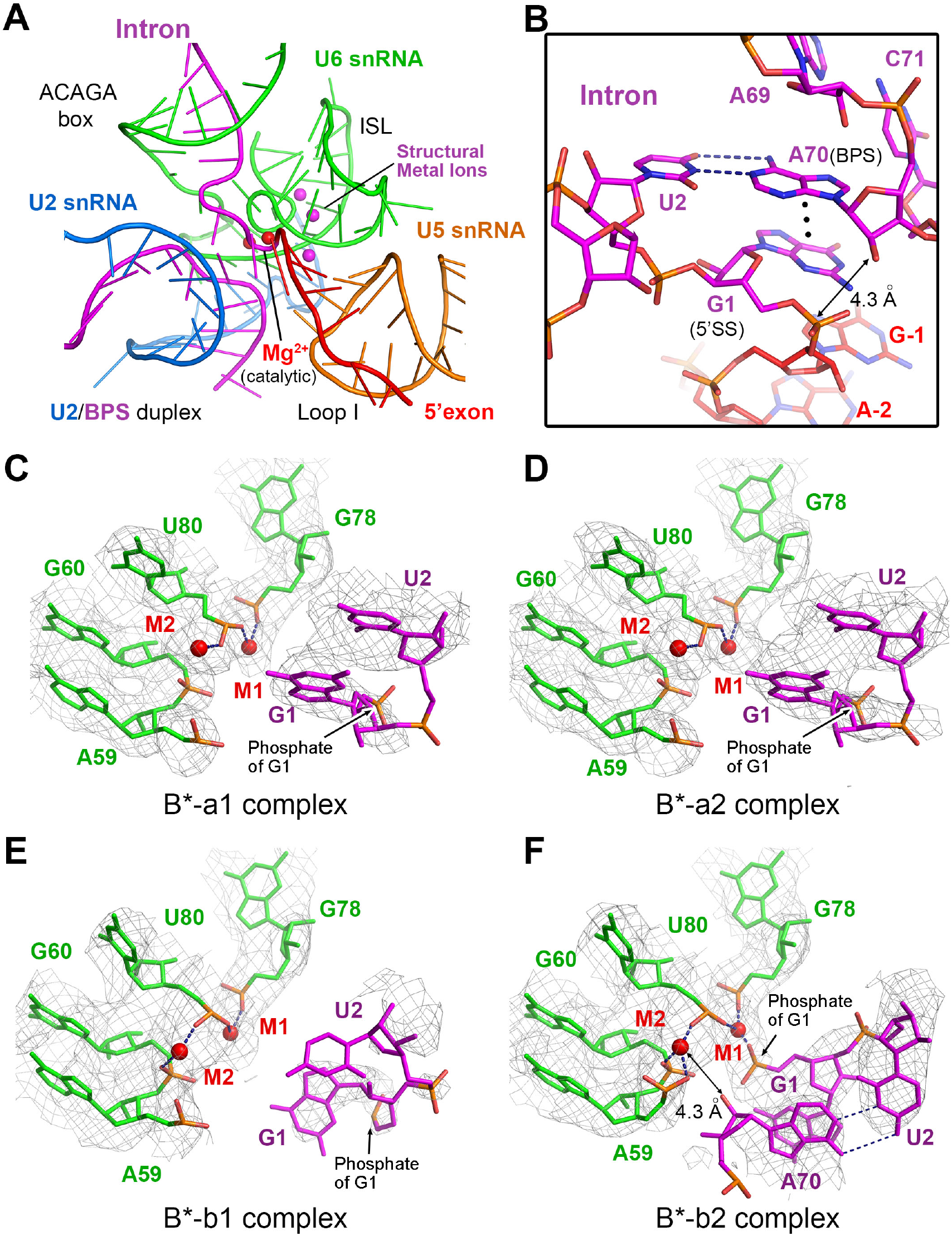
The splicing active site in the B^*^-b2 complex and the coordination of catalytic metal ions in the B^*^ complexes. (A) Structure of the splicing active site in the B^*^-b2 complex. The catalytic center comprises the intramolecular stem loop (ISL) of U6 snRNA and the associated Mg^2+^ ions, helix I of the U2/U6 duplex, and loop I of U5 snRNA. The catalytic and structural metal ions are shown as red and magenta spheres, respectively. The U2/BPS duplex is positioned in close proximity to the 5’SS in the active site. (B) A close-up view on the recognition of the invariant adenine nucleotide A_70_ of the BPS at the active site center of the B^*^-b2 complex. A_70_ of the BPS is recognized by the conserved dinucleotide G_1_U_2_ at the 5’-end of the 5’SS, through A_70_-U_2_ base-pairing and A_70_-G_1_ base stacking. The nucleophile 2’-OH of A_70_ is located only ~4.3 Å away from the phosphate of G_1_, nearly ready for nucleophilic attack. (C) A close-up view on the coordination of the catalytic metals M1 and M2 in the B^*^-a1 complex. M2 is coordinated by a phosphate from U_80_ of U6 snRNA. M1 is bound by the phosphate oxygens of G_78_ and U_80_ from U6 snRNA. (D) A close-up view on the coordination of M1 and M2 in the B^*^-a2 complex, which is identical to that in the B^*^-a1 complex. (E) A close-up view on the coordination of M1 and M2 in the B^*^-b1 complex. Compared to that in B^*^-a1 or -a2, M2 is coordinated by an additional phosphate from G_60_ of U6 snRNA. The coordination of M1 is identical to that in the B^*^-a1 and -a2 complexes. (F) A close-up view on the coordination of M1 and M2 in the B^*^-b2 complex. M2 is coordinated by three phosphates from A_59_, G_60_ and U_80_ of U6 snRNA. But the 2’-OH of A_70_ is yet to be activated by M2 in this complex, with a distance of approximately 4.3 Å. M1 stabilizes the leaving group, the phosphate of G_1_ in the 5’SS, through direct interaction. M1 is also coordinated by the phosphate oxygens of G_78_ and U_80_ from U6 snRNA.

**Figure 4.**
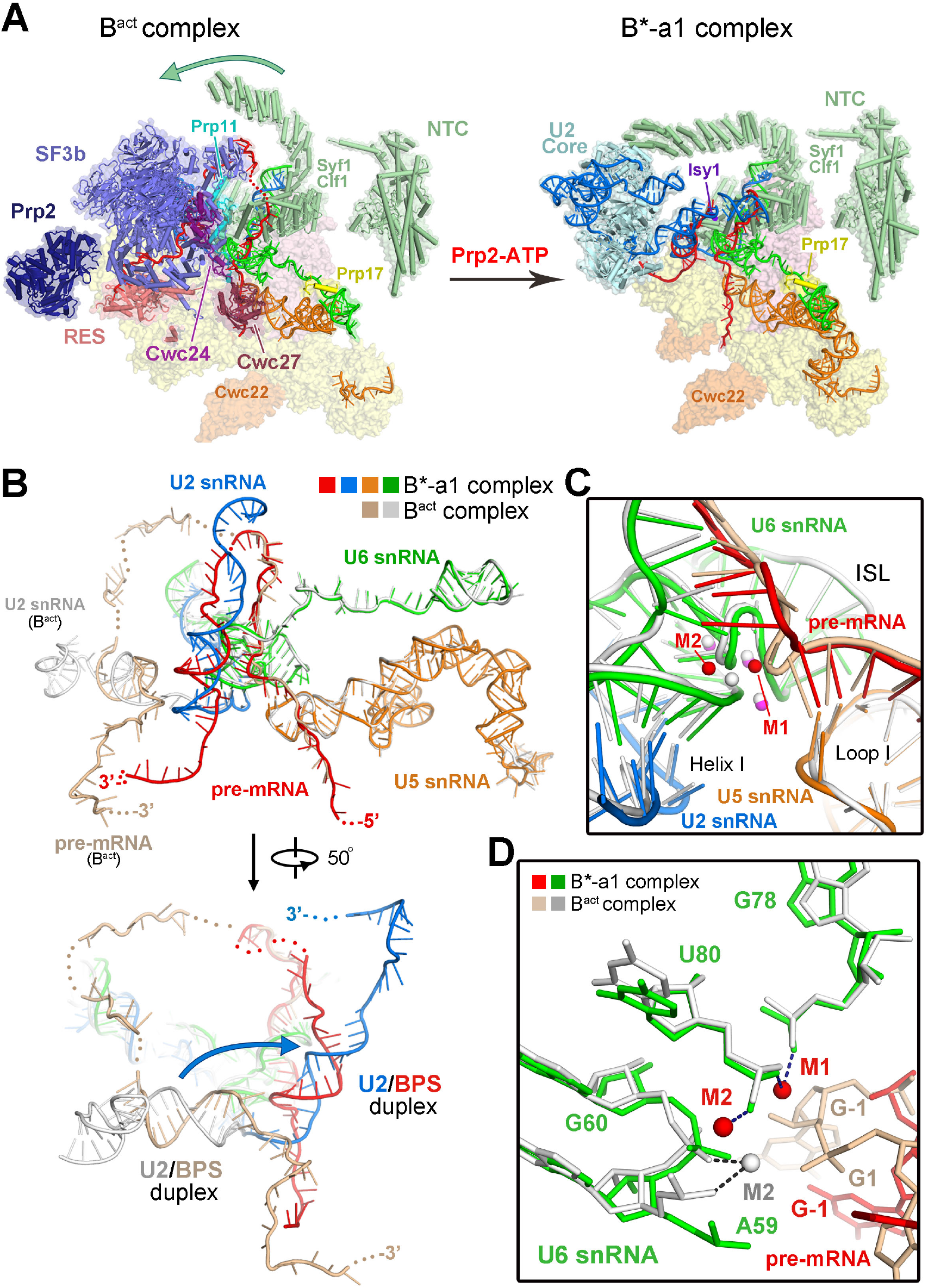
Structural changes of the B^act^-to-B^*^ transition in *S. cerevisiae.* (A) Structural changes of the protein components during the transition from the B^act^ to B^*^-a1 complex. In the B^act^ complex (Yan et al., 2016), the SF3b complex and the RES complex recognize the U2/BPS duplex and the downstream RNA sequences of the intron, respectively. Upon Prp2-mediated remodeling, SF3a, SF3b, RES and the splicing factors Cwc24 and Cwc27 are dissociated to release the U2/BPS duplex and the 5’SS. The U2 snRNP core and the NTC proteins Syf1 and Clf1 undergo marked translocation. Isy1 associates with the RNA elements at the active site center. Consequently, the U2/BPS duplex is delivered close to the active site. (B) Structural comparison of the overall RNA elements between the B^act^ complex (Yan et al., 2016) and the B^*^-a1 complex. Two views are shown. The RNA elements in the B^*^-a1 complex are colored identically as those in Figure 2A; in the B^act^ complex, pre-mRNA is colored wheat and all other RNA elements are shown in gray. During the B^act^-to-B^*^ transition, the U2/BPS duplex undergoes a drastic translocation to the vicinity of the active site. U5 and U6 snRNAs and the first 30 nucleotides of U2 snRNA remain unchanged. (C) Structural comparison of the active site between the B^act^ (Yan et al., 2016) and B^*^-a1 complexes. Compared to the B^*^-a1 complex, M1 is yet to be loaded in the B^act^ complex, M2 undergoes a positional shift. (D) Comparison on the coordination of the catalytic metals between the B^act^ and B^*^-a1 complexes. In the B^act^ complex, M1 is yet to be loaded, M2 is coordinated by two phosphates from A_59_ and G_60_ of U6 snRNA. In the B^*^-a1 complex, M1 is already loaded into the active site and coordinated by two phosphates from G_78_ and U_80_ of U6 snRNA. M2 is coordinated by a phosphate from U_80_ of U6 snRNA.

Concurrent with the flux of the protein components, some of the RNA elements also undergo drastic structural changes. Compared to the B^act^ complex, the U2 snRNA sequences downstream of nucleotide 30 and the pre-mRNA downstream of the 5’SS are drastically translocated in the B^*^-a1 complex (Figure 4B). The interaction between stem IIb of U2 snRNA and the C-terminal domain of Ecm2 is clearly present in the four B^*^ complexes (Figure S5F). In addition, stem IIc of U2 snRNA, which is thought to promote the 5’SS cleavage (Hilliker et al., 2007; Perriman and Ares, 2007), is formed during the transition from the B^act^ to B^*^-a1 complex (Figure S7E). The U2/BPS duplex, which was protected by the SF3b complex and located ~50 Å away from the active site in the B^act^ complex (Yan et al., 2016), undergoes a movement of 30-40 Å into the vicinity of the 5’SS in the B^*^-a1 complex. In contrast, U5 and U6 snRNAs and the first 30 nucleotides of U2 snRNA remain unchanged during the transition (Figure 4B). Consequently, the RNA elements in the active site remain nearly identical between the B^act^ and B^*^ complexes (Figure 4C). The only notable difference is that, compared to the B^*^-a1 complex, M1 is yet to be loaded in the B^act^ complex (Yan et al., 2016) (Figure 4D; Figure S6C, left panel).

In the B^act^ complex (Yan et al., 2016), the RNA elements at the active site (including pre-mRNA and snRNAs) are coated by protein components and stabilized by Prp11, Cwc24 and the 1585-loop (also known as α-finger) of Prp8, which allow little movement of the RNA elements. Consequently, the M1 metal is not loaded, because the phosphate oxygens from G78 and U80 of U6 snRNA are not in the right positions to coordinate M1. In the B^*^ complex, most of the coated proteins have been dissociated through the action of Prp2. The RNA elements at the active site are less constrained; the ISL of U6 snRNA undergoes slight conformational adjustment to allow coordination of M1 by the phosphate oxygens from G78 and U80.

### Structural changes of the B^*^-to-C transition

Branching occurs in the B^*^ complex, resulting in the C complex. Unlike the B^act^-to-B^*^ transition, the B^*^-to-C transition has been predicted to involve few structural changes (Shi, 2017b; Wan et al., 2016b). Indeed, the snRNA elements and much of the pre-mRNA are nearly identical between the B^*^-b2 and C complexes (Wan et al., 2016b) (Figure 5A). The RNA elements in the active site of the B^*^-b2 complex can be superimposed to those of the C complex with a root-mean-squared deviation of 0.46 Å for 80 nucleotides (Figure 5B). The only apparent difference is that the 3’-5’ covalent linkage between the 5’-exon and 5’SS in the B^*^-b2 complex is broken and replaced by the 2’-5’ phosphodiester bond between A_70_ of the BPS and G_1_ of the 5’SS in the C complex (Wan et al., 2016b) (Figure 5C). As a consequence of branching, coordination of the catalytic metals in the C complex is different from that in the B^*^-b2 complex (Figure 5D; Figure S6C, middle and right panels). In the C complex, M1 is coordinated in a planar fashion by four ligands: 3’-OH of G_-1_ of the 5’-exon and three phosphate oxygens from G_78_ and U_80_ of U6 snRNA and G_1_ of 5’SS (Wan et al., 2016b). In the B^*^-b2 complex, M1 is only coordinated by three ligands but does not interact with 3’-OH of G_-1_ of the 5’-exon. Notably, both M2 and the phosphates of G_-1_ and G_1_ undergo apparent positional shifts in the B^*^-to-C transition (Figure 5D). In the C complex, M2 is bound by two phosphate oxygens from A_59_ and U_80_ of U6 snRNA (Wan et al., 2016b); in the B^*^-b2 complex, M2 is coordinated by three phosphates from A_59_, G_60_ and U_80_ of U6 snRNA.

**Figure 5.**
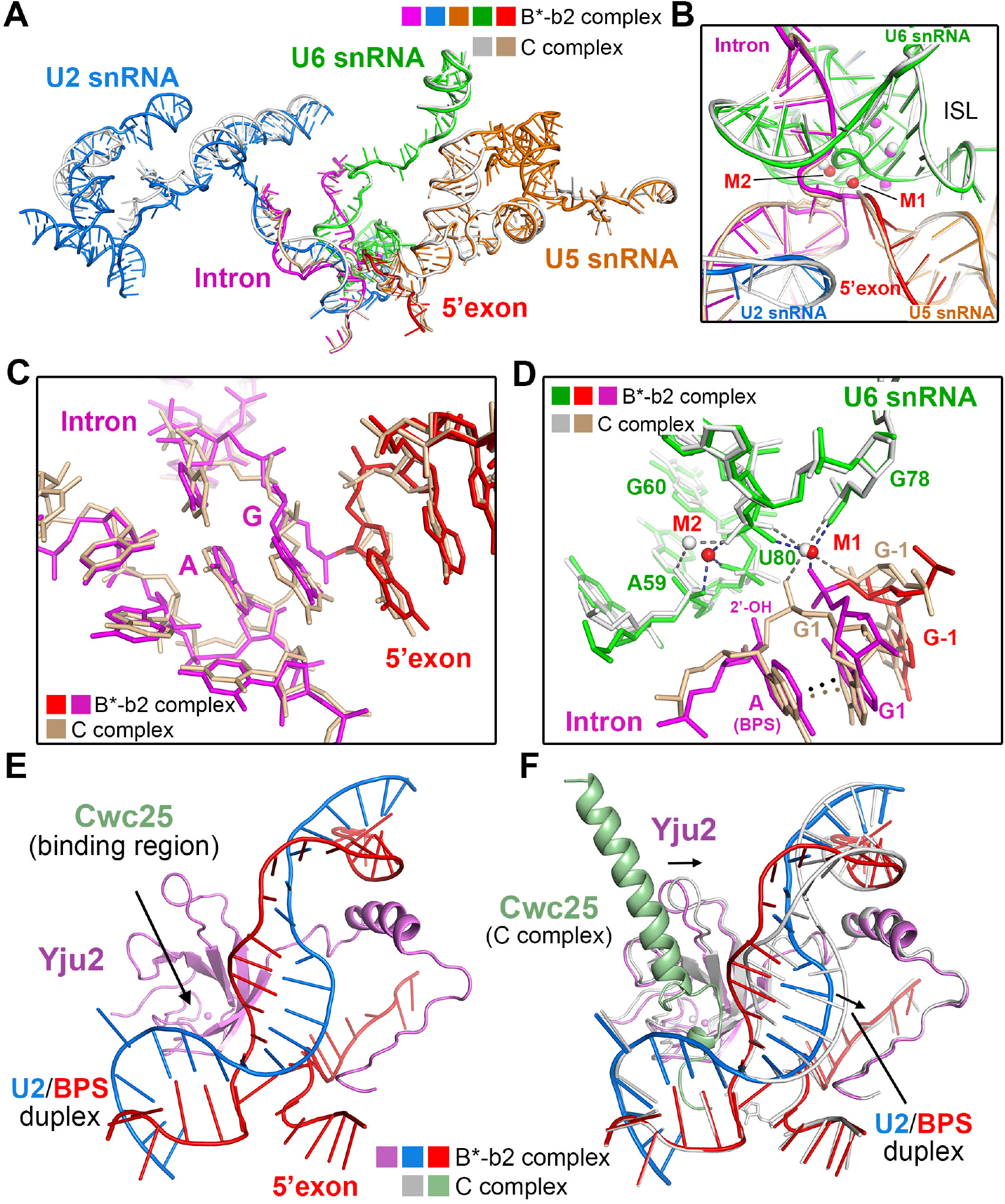
Structural comparison between the B^*^-b2 complex and the C complex from *S. cerevisiae.* (A) Overall structural comparison of the RNA elements between the B^*^-b2 complex and the C complex (Wan et al., 2016b). The RNA elements in the B^*^-b2 complex are colored identically as those in Figure 2A; in the C complex, pre-mRNA is colored wheat and all other RNA elements are shown in gray. Except for minor differences in the 3’-end region of U2 snRNA and the BPS region, structures of all other RNA elements are nearly identical. (B) A close-up view on the structural comparison of the active site between the B^*^-b2 and C complexes. (C) A close-up view on the structural comparison of the region surrounding A_70_ of the BPS and G_1_ of the 5’SS between the B^*^-b2 and C complexes. The nucleophile and the acceptor are positioned close to each other in the B^*^-b2 complex but are covalently linked together in the C complex. (D) A close-up view on the structural comparison of catalytic metal coordination between the B^*^-b2 and C complexes. In the C complex, M1 is coordinated in a planar fashion by four ligands: 3’-OH of G_-1_ of the 5’-exon and three phosphate oxygens from G_78_ and U_80_ of U6 snRNA and G_1_ of 5’SS. In the B^*^-b2 complex, M1 is only coordinated by three ligands and no longer interacts with 3’-OH of G_-1_ of the 5’-exon. In the C complex, M2 is bound by two phosphate oxygens from A_59_ and U_80_ of U6 snRNA; in the B^*^-b2 complex, M2 is coordinated by three phosphates from A_59_, G_60_ and U_80_ of U6 snRNA. (E) A close-up view on the region occupied by the step I splicing factors in the B^*^-b2 complex. The N-terminal loop and downstream β-sheet domain of Yju2 bind to the 5’SS and the U2/BPS duplex, respectively. Cwc25 remains to be recruited into the active site. (F) A close-up view on the region occupied by the step I splicing factors between the B^*^-b2 complex and the C complex. The β-sheet domain of Yju2 and the U2/BPS duplex exhibit apparent positional shifts between the B^*^-b2 and C complexes.

In the B^*^-b2 complex, the step I splicing factor Yju2 is already loaded into the active site, with its N-terminal loop and β-sheet domain binding to 5’SS and U2/BPS duplex, respectively (Figure 5E). Consistent with previous biochemical analysis (Liu et al., 2007; Ohrt et al., 2012; Warkocki et al., 2009), the presence of Yju2 in the structure suggests its recruitment prior to or during the formation of the B^act^ complex because no recombinant Yju2 is required in our preparation of the B^*^ complex. However, the other step I factor Cwc25, which is thought to be required after Yju2 to promote branching (Chiu et al., 2009; Warkocki et al., 2009), is still absent in the B^*^-b2 complex. A close-up comparison of the region occupied by the step I factors between the B^*^-b2 and C complexes reveals apparent positional shifts for the β-sheet domain of Yju2 and the U2/BPS duplex (Figure 5F). These differences are accompanied by conformational changes of the U2/BPS duplex.

Notably, Cwc25 is the last protein factor recruited to the spliceosome before branching and the recruitment of Cwc25 is thought to require Yju2 (Chiu et al., 2009). Consistent with this conclusion, the active site conformation in the B^*^-b2 complex, which is stabilized by Yju2, is conducive for Cwc25 recruitment (Figure 5E). The recruitment of Cwc25, in turn, likely leads to a positional shift of the Yju2 β-sheet domain and movement of U2/BPS towards the active site center (Figure 5F).

## Discussion

Spliceosome remodeling from the B^act^ to the B^*^ complex is driven by Prp2 and its cofactor Spp2 (Kim and Lin, 1993; King and Beggs, 1990; Roy et al., 1995). SF3a, SF3b, RES and the splicing factors Cwc24 and Cwc27 are dissociated from the B^act^ complex (Lardelli et al., 2010; Ohrt et al., 2012), and the step I splicing factors Yju2 and Cwc25 are required for branching after the action of Prp2 (Chiu et al., 2009; Krishnan et al., 2013; Liu et al., 2007). Yju2 by itself is insufficient to yield the intron lariat-3’-exon intermediate (Liu et al., 2007; Warkocki et al., 2009). Branching is dramatically stimulated by the addition of Cwc25, suggesting its indispensable role in promoting efficient branching (Krishnan et al., 2013; Warkocki et al., 2009). Despite these biochemical studies, how Yju2 and Cwc25 differentially facilitate the branching reaction remains unclear. Our structural analysis provides a plausible answer to this key question.

Structure determination of the B^*^ complex allows proposition of the structural changes at the active site during the B^act^-to-B^*^-to-C transition in *S. cerevisiae* (Figure 6A). This transition is triggered by Prp2, which may pull the 3’-end of the intron (Liu and Cheng, 2012; Yan et al., 2016), dissociating proteins that are associated with the BPS and 5’SS and allowing translocation of the U2/BPS duplex into the active site (Lardelli et al., 2010; Ohrt et al., 2012; Warkocki et al., 2009; Warkocki et al., 2015). The nucleophile-containing A_70_ of the BPS is stabilized through base-pairing interaction with U_2_ and base-stacking interaction with G_1_ at the 5’-end of the 5’SS. This conformation is supported by Isy1 and Yju2. Upon recruitment of Yju2 and Cwc25 into the active site, the nucleophile from U2/BPS duplex is moved into the close proximity of the acceptor in the 5’SS, forming the B^*^ complex. Branching occurs instantaneously, resulting in a covalent linkage between the 2’-OH of BPS A_70_ and the 5’-phosphate of 5’SS G_1_ in the C complex (Galej et al., 2016; Wan et al., 2016b).

**Figure 6.**
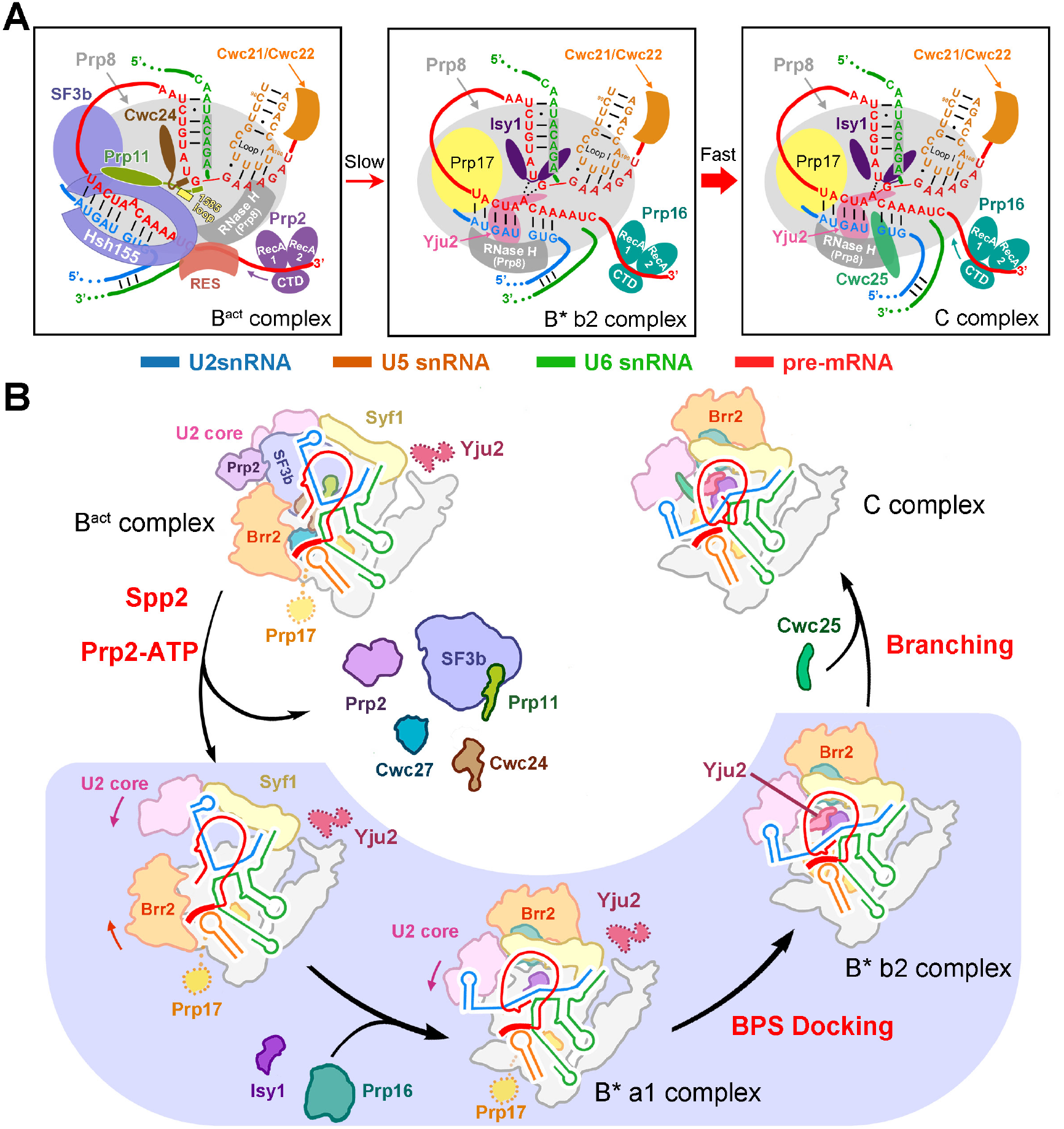
A mechanistic model of branching catalyzed by the spliceosome in *S. cerevisiae.* (A) A cartoon diagram of the structural changes at the active site during the B^act^-to-B^*^-to-C transition. In this model, Prp2 may pull the 3’-end of the intron, dissociating proteins that are associated with the BPS and 5’SS and allowing the translocation of the U2/BPS duplex into the active site. The nucleophile-containing adenine nucleobase in BPS is stabilized through base-pairing with U_2_ and base-stacking with G_1_ at the 5’-end of the 5’SS. Upon binding of the step I splicing factor Cwc25, the U2/BPS duplex is pushed closer to the 5’SS, allowing the branching to occur and resulting in the C complex. (B) A schematic diagram of spliceosome remodeling by Prp2 and the branching reaction catalyzed by the spliceosome in *S. cerevisiae.* During the B^act^-to-B^*^ transition, the ATPase/helicase Prp2 and its cofactor Spp2 mediate the dissociation of the SF3a and SF3b complexes and the splicing factors Cwc24 and Cwc27, freeing the 5’SS and the U2/BPS duplex. The reorganization of the protein components may allow translocation of the U2/BPS duplex into the vicinity of the splicing active site, forming a partially catalytically activated complex (bottom middle), exemplified by the B^*^-a1 complex. Next, Yju2 is loaded to the spliceosome catalytic center, pushing the U2/BPS duplex closer to the 5’SS (bottom right), forming the B^*^-b2 complex. In the final step, Cwc25 is recruited, allowing fine adjustment of the active RNA and protein elements and consequent branching reaction. The resulting spliceosome is the C complex.

On the basis of previous biochemical evidence and our structural observations, we further propose that the B^act^-to-B^*^ transition may occur in several distinct steps (Figure 6B). First, the ATPase/helicase Prp2 and its cofactor Spp2 mediate the dissociation of the SF3a and SF3b complexes and the splicing factors Cwc24 and Cwc27, freeing the 5’SS and the U2/BPS duplex (Lardelli et al., 2010; Ohrt et al., 2012; Roy et al., 1995; Warkocki et al., 2009). In the resulting intermediary spliceosome (Figure 6B, bottom left), a number of the components and subcomplexes, which include Brr2, U2 snRNP core and two NTC proteins Syf1 and Clf1 of NTC, may exhibit dynamic conformation. Next, recruitment of Isy1 and the WD40 domain of Prp17 in the active site may lead to reorganization of the surrounding components, allowing translocation of the U2/BPS duplex into the approximately 20-Å range of the 5’SS and forming a partially catalytically activated complex (Figure 6B, bottom middle). This spliceosome may exhibit a range of dynamic conformations, exemplified by the a1 and a2 conformations of the *ACT1* B^*^ complex and the b1 conformation of the *UBC4* B^*^ complex prior to Yju2 loading. Then, the recruitment of Yju2 further reorganizes the active site, pushing the U2/BPS duplex into the close proximity of the 5’SS as observed in the B^*^-b2 complex (Figure 6B, bottom right). Finally, Cwc25 is recruited, allowing fine adjustment of the active site elements and consequent execution of the branching reaction (Wan et al., 2016b). The resulting spliceosome is the C complex.

Structure of the Yju2-containing B^*^ complex reveals, for the first time, the interaction between the highly conserved dinucleotide G_1_U_2_ at the 5’-end of the 5’SS and the invariant A_70_ of the BPS just before branching (Figure 3B). In particular, the nucleophile-containing A_70_ is recognized and partially stabilized by U_2_ through base-pairing interactions. Replacement of U_2_ by any other nucleotide may weaken or disrupt the interaction thus reducing the splicing level (Aebi et al., 1986; Aebi et al., 1987; Lamond et al., 1987). This analysis suggests that formation of these interactions may represent a potential check-point for proofreading the first transesterifi cation. A similar check-point may exist just prior to exon ligation, whereby the invariant G_1_ of the 5’SS and A_70_ of the BPS recognize the dinucleotide AG of the 3’SS through base-paring and stacking interactions as observed in the P complex (Bai et al., 2017; Liu et al., 2017; Wilkinson et al., 2017) (Figure 7A). Utilization of such conserved nucleotides towards specific interactions may constitute an additional layer of regulation to safeguard the splicing specificity and efficiency.

**Figure 7.**
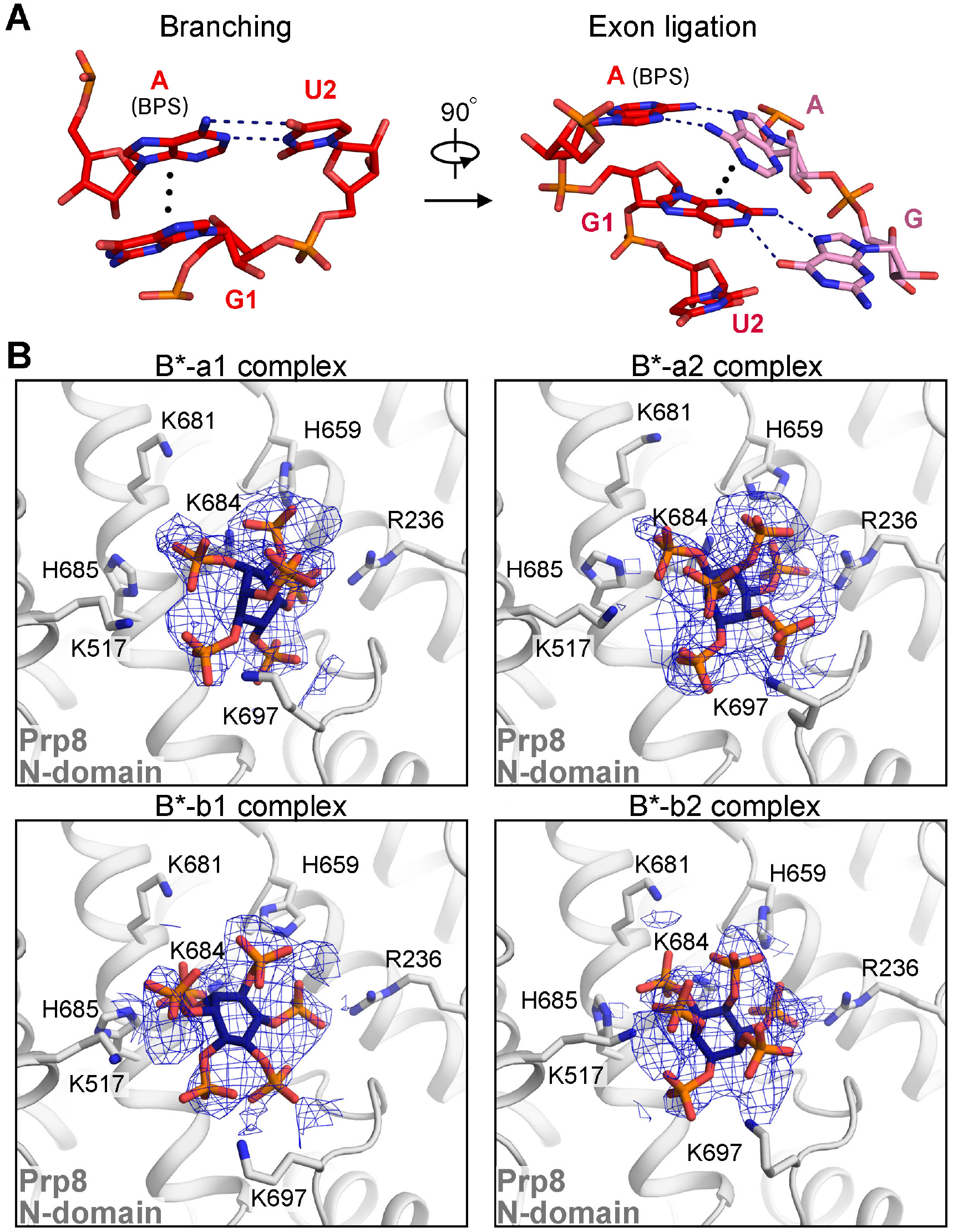
Conserved structural features in the B^*^ complex and other functional states of the assembled spliceosome. (A) Interactions among conserved intron sequences in branching and exon ligation. In the B^*^ complex preceding the branching reaction (left panel), the nucleophile-containing A_70_ of the BPS is recognized through base-pairing with U_2_ and base-stacking against G_1_ of the 5’SS. In the P complex following exon ligation (right panel), the dinucleotides AG at the 3’-end of the 3’-splice site (3’SS) are stabilized by the invariant A_70_ of the BPS and G_1_ of the 5’SS through base-pairing and base-stacking interactions, respectively. (B) A small molecule interacts with Prp8 in all four B^*^ complexes. This small molecule, identified as inositol hexaphosphate (IP6), is identically present in the yeast B^act^ through ILS complexes. The EM density maps of IP6 and its interactions with Prp8 in the four distinct conformations of the B^*^ complex are shown here. IP6 is located in a positively charged cavity formed in the N-domain of Prp8.

Previous studies reported the proofreading function of Prp16 and Prp22 before branching and exon ligation, respectively (Burgess and Guthrie, 1993; Koodathingal et al., 2010; Mayas et al., 2006; Query and Konarska, 2004; Semlow et al., 2016; Villa and Guthrie, 2005). In the structure of the four B^*^ complexes, the binding site for Prp16 is exposed. But the stable association of Prp16 requires the presence of both Cwc25 and Yju2, as observed in the structure of the C complex (Galej et al., 2016; Wan et al., 2016b). Consistently, in our structures of the B^*^ complex, the EM density for Prp16 is insufficient for its conclusive assignment.

The overall structures of the *ACT1* B^*^ complex and the *UBC4* B^*^ complex are unsurprisingly similar, because the two pre-mRNA molecules share exactly the same sequences in the three signature elements: 5’SS, BPS, and 3’SS. Consequently, the 5’SS is recognized identically by the ACAGA box of U6 snRNA in either *ACT1* or *UBC4* B^*^ complex (Figure 2D,E, upper panels). The BPS also forms the same duplex with the conserved sequences of U2 snRNA in either B^*^ complex (Figure 2B,C). Such structural similarities even extend to a small molecule, which has been identified as inositol hexaphosphate (IP6). This small molecule is identically bound to the N-domain of Prp8 not just in all four structures of the B^*^ complex (Figure 7B) but perhaps more importantly in all yeast spliceosomes between the functional states of B^act^ and ILS. The function of this small molecule in splicing remains to be investigated.

The *ACT1* B^*^ complex and the *UBC4* B^*^ complex differ in the 3’-end nucleotides of their 5’-exons. These differences determine the strengths of interactions with loop I of U5 snRNA. Only a single nucleotide from the 5’-exon of *ACT1* base-pairs with loop I, whereas four nucleotides from the 5’-exon of *UBC4* form a duplex with loop I (Figure 2D,E, lower panels). The sequence variation in *ACT1* versus *UBC4* also results in differences in overall structure of the spliceosome (Figure 1), conformation of key RNA elements (Figure 2), and coordination of catalytic metal ions (Figure 3). These structurally documented differences constitute compelling evidence for substrate-specific conformations of the spliceosome in a major functional state - in this case the B^*^ complex. These structural observations are consistent with the biochemical finding that depletion or mutation of spliceosomal core components exhibits differential effects on splicing substrates (Campion et al., 2010; Clark et al., 2002; Kawashima et al., 2009; Park et al., 2004; Pleiss et al., 2007; Saltzman et al., 2011). We further speculate that substrate-specific conformation may be a general phenomenon for all major functional states of the spliceosome.

The B^*^ complex, if simultaneously containing Cwc25 and Yju2, would instantaneously proceed to branching and is thus extremely transient. To obtain the B^*^ complex, we took the *in vitro* assembly approach using recombinant Prp2 and Spp2 (Bao et al., 2017; Warkocki et al., 2009). This approach allowed us to observe four distinct conformations of the B^*^ complex assembled on two different pre-mRNA. Yju2 is only present in one of the *UBC4* B^*^ conformers. These four structures might represent the B^*^ complex just preceding the spliceosomal state that catalyzes branching. Three structures show that, in the absence of Cwc25 and Yju2, the U2/BPS duplex cannot be placed into the correct location for the branching reaction, explaining the essential function of the step I splicing factors (Chiu et al., 2009; Krishnan et al., 2013; Liu et al., 2007; Warkocki et al., 2009). The recruitment of Yju2 into the active site allows the U2/BPS duplex to move into the close proximity of the 5’SS; but in the absence of Cwc25, the nucleophile remains to be activated by the M2 metal and the overall active site conformation is subtly unsuitable for the branching reaction.

## STAR METHODS

Detailed methods are provided in the online version of this paper and include the following:

- KEY RESOURCES TABLE
- CONTACT FOR REAGENT AND RESOURCE SHARDING
- EXPERIMENTAL MODEL AND SUBJECT DETAILS

- Cell lines
- METHOD DETAILS

- CBP tagging of in *S. cerevisiae*
- Preparation of the yeast whole cell extract
- Expression and purification of Prp2 and Spp2
- Preparation of the pre-mRNA
- Assembly and purification of the yeast B^act^ (ΔPrp2) complex
- Remodeling of the B^act^ (ΔPrp2) complex by Prp2 and Spp2
- EM data acquisition
- Data processing
- Model building and refinement
- QUANTIFICATION AND STATISTICAL ANALYSIS
- DATA AND SOFTWARE AVAILABILITY

- Data Resources

## SUPPLEMENTAL INFORMATION

Supplemental Information includes three tables and seven figures and can be found with this article online at XXX.

### Author Contributions

R.B. prepared the whole cell extract, pre-mRNA substrates, and recombinant proteins Prp2 and Spp2. R.W. and R.B. purified the yeast spliceosomes and prepared the cryo-EM samples. R.W., R.B., and J.L. collected and processed the EM data. R.W. calculated the EM map and built the atomic model. C.Y. gave advices on data processing and model building. All authors contributed to structure analysis. R.W. and R.B. contributed to manuscript preparation. Y.S. designed and guided the project and wrote the manuscript.

## ACKNOWLEDGMENTS

We thank the Tsinghua University Branch of China National Center for Protein Sciences (Beijing) for providing the facility support. The computation was completed on the “Explorer 100” cluster system of Tsinghua National Laboratory for Information Science and Technology. This work was supported by funds from the National Natural Science Foundation of China (31621092 and 31430020), the National Postdoctoral Program for Innovative Talents (BX201800125 to R.W.) and the Ministry of Science and Technology (2016YFA0501100 to J.L.) The atomic coordinates have been deposited in the Protein Data Bank with the following accession codes XXXX. The EM maps have been deposited in the EMDB with the following accession codes YYYY. The authors declare no competing financial interests. Correspondence and requests for materials should be addressed to Y. Shi.

