## Supplementary Figures, Methods, Tables S1-S3 for "Structures of the Catalytically Activated Yeast Spliceosome Reveal the Mechanism of Branching"

#### Supplemental Figures

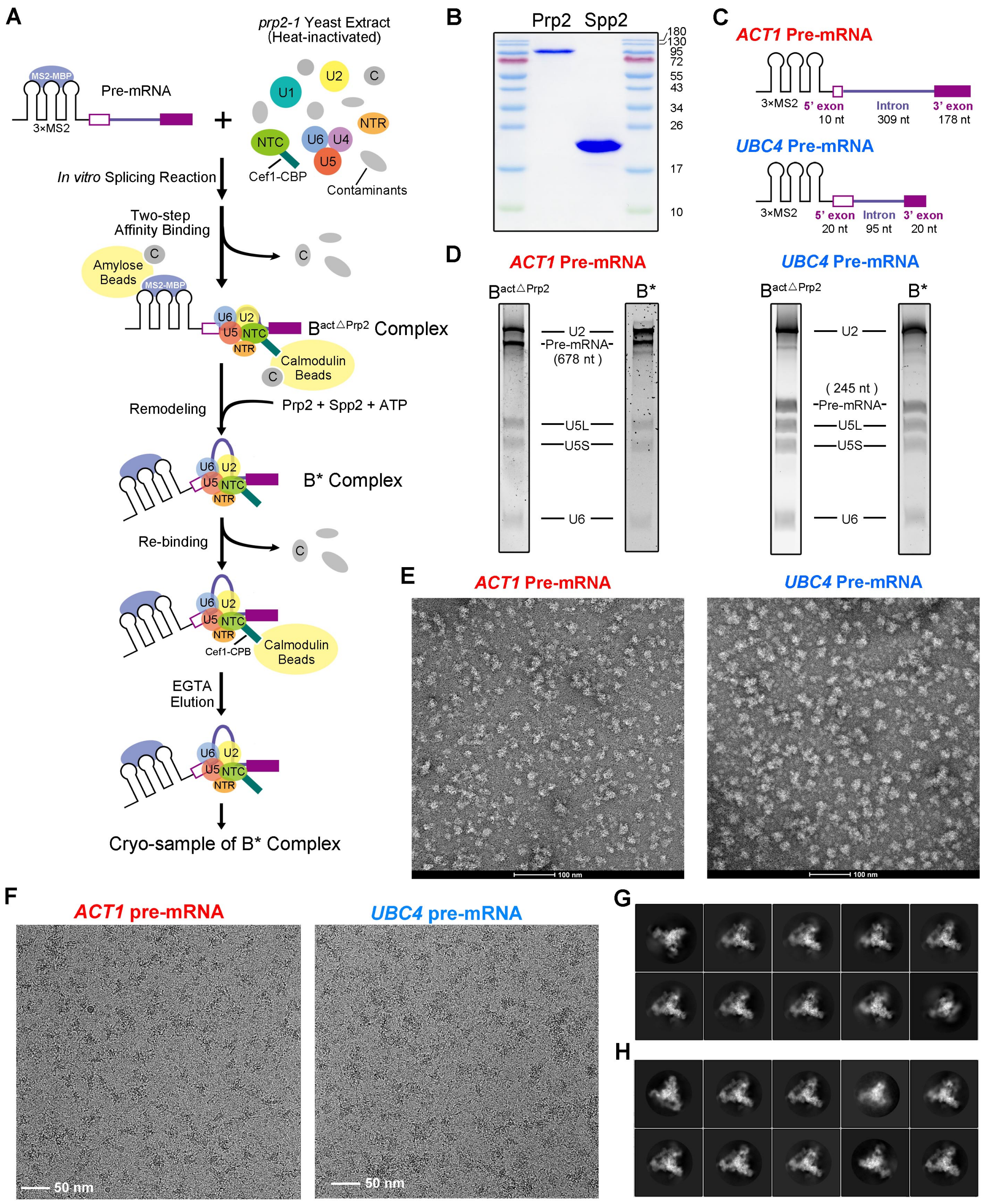


**Figure S1 Purification and EM analysis of the spliceosomal B$^*$ complexes from *S. cerevisiae*. Related to Figure 1.** (A) A schematic diagram of the purification protocol for the B$^*$ complex. (B) The purified recombinant proteins Prp2 and Spp2 are shown on a Coomassie-stained SDS-PAGE gel. (C) A schematic diagram of the two pre-mRNA substrates used in this study. (D) Analysis of the B$^*$ complex assembled on the *ACT1* pre-mRNA and the *UBC4* pre-mRNA eluted from the affinity column. Shown here are representative results of a urea PAGE gel. All five snRNAs are clearly present. (E) A representative negative-staining EM micrograph of the purified *ACT1* $B^*$ complex and *UBC4* $B^*$ complex. Scale bar, 100 nm. (F) A representative cryo-EM micrograph of the purified *ACT1* $B^*$ complex and *UBC4* $B^*$ complex. Scale bar, 50 nm. (G) Representative two-dimensional (2D) class averages of the electron micrographs for the *ACT1* $B^*$ complex. (H) Representative 2D class averages of the electron micrographs for the *UBC4* $B^*$ complex.

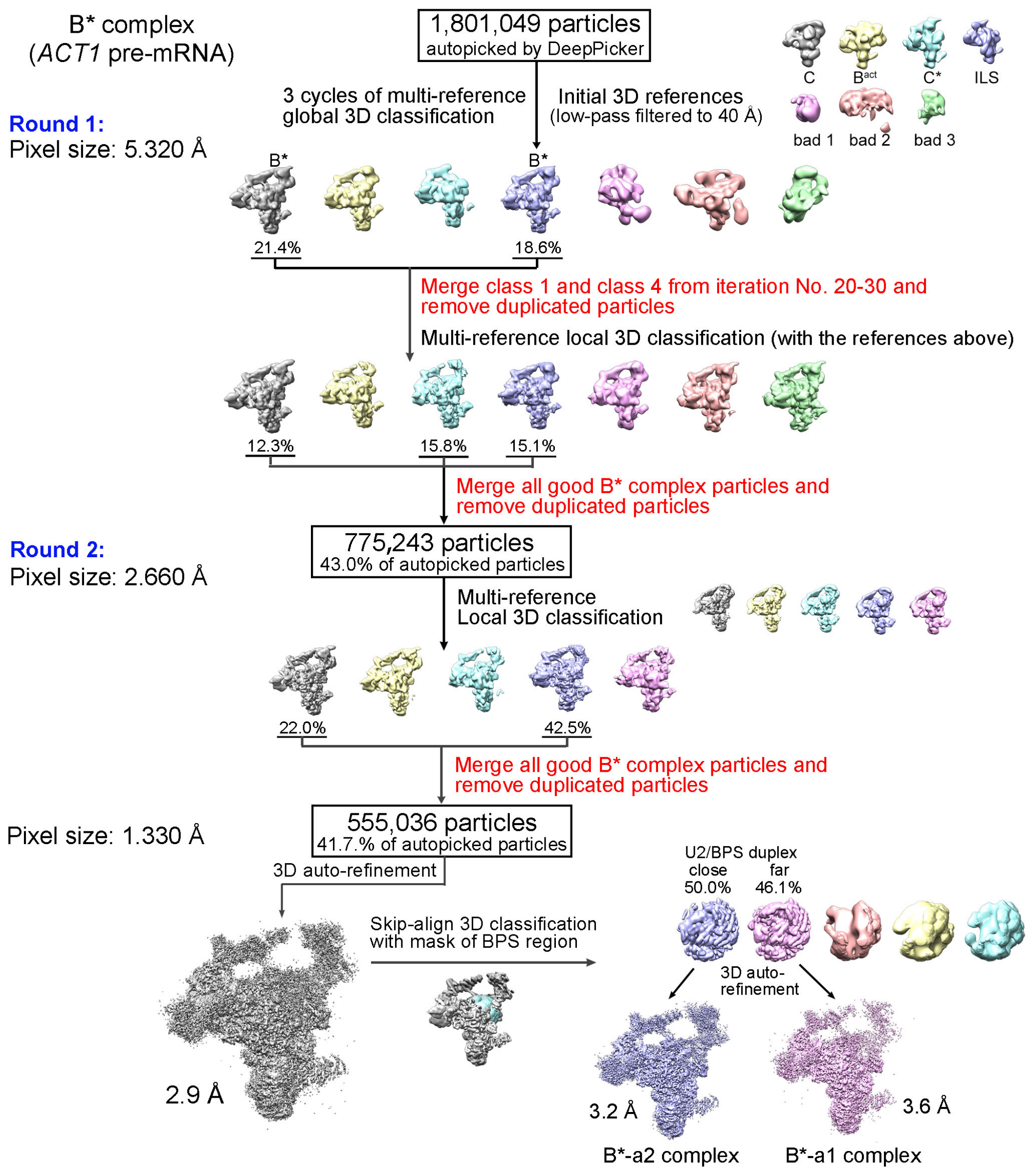


**Figure S2 A flow chart for the cryo-EM data processing and structure determination of the spliceosomal B$^*$ complex assembled on the *ACT1* pre-mRNA from *S. cerevisiae*. Related to Figures 1 & 2** On the basis of the FSC value of 0.143, the final reconstruction has an average resolution of 2.9 Å. Two distinct conformational states of the *ACT1* B$^*$ complex were further differentiated, one at 3.2 Å resolution (B$^*$-a2) and the other at 3.6 Å (B$^*$-a1). Please refer to Materials and Methods for details.

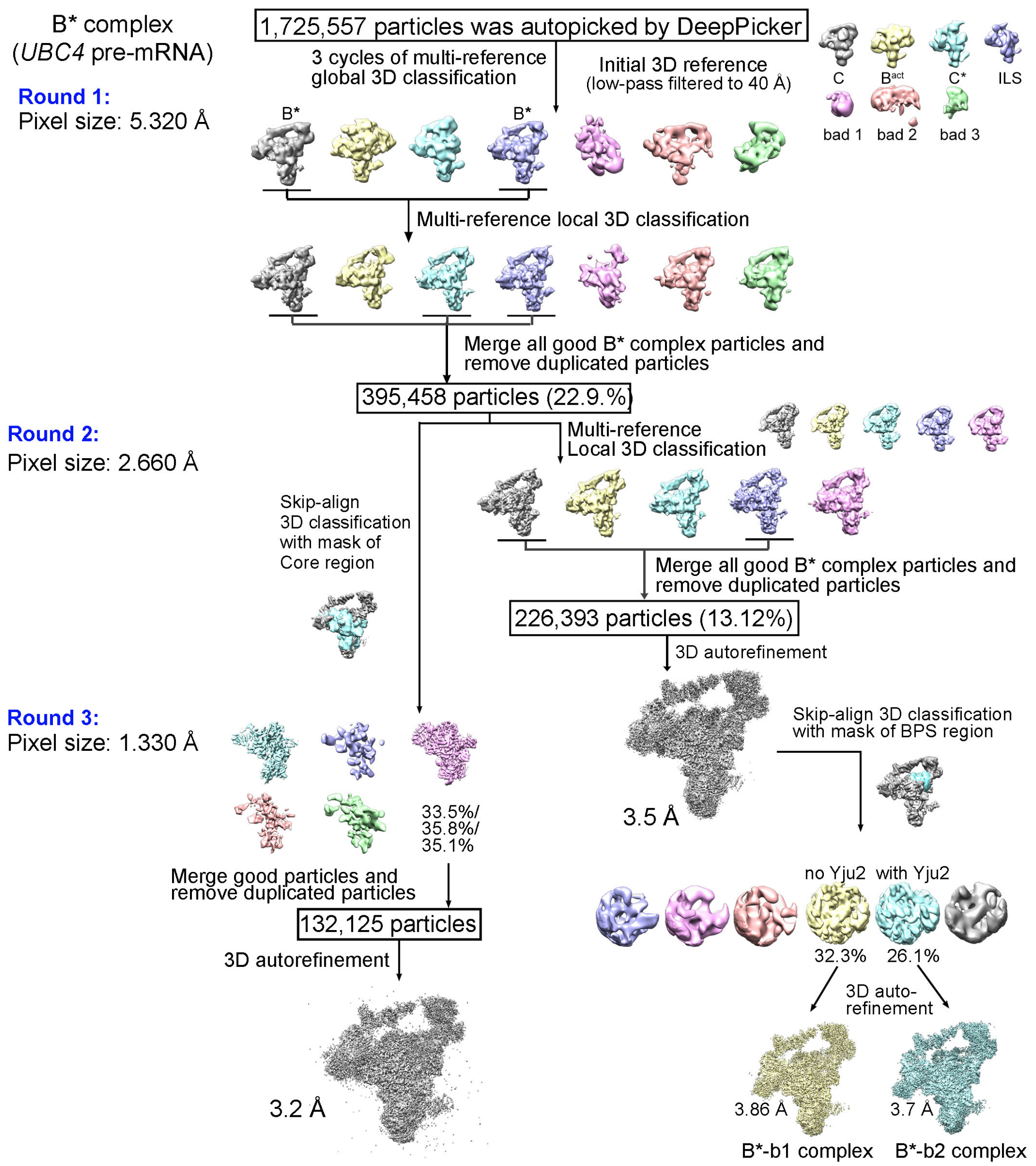


**Figure S3 A flow chart for the cryo-EM data processing and structure determination of the spliceosomal B$^*$ complex assembled on the *UBC4* pre-mRNA from *S. cerevisiae*. Related to Figures 1 & 2** On the basis of the FSC value of 0.143, the final reconstruction has an average resolution of 3.2 Å. Two distinct conformational states of the *UBC4* B$^*$ complex were further differentiated, one at 3.86 Å resolution (B$^*$-b1) and the other at 3.7 Å (B$^*$-b2). Please refer to Materials and Methods for details.

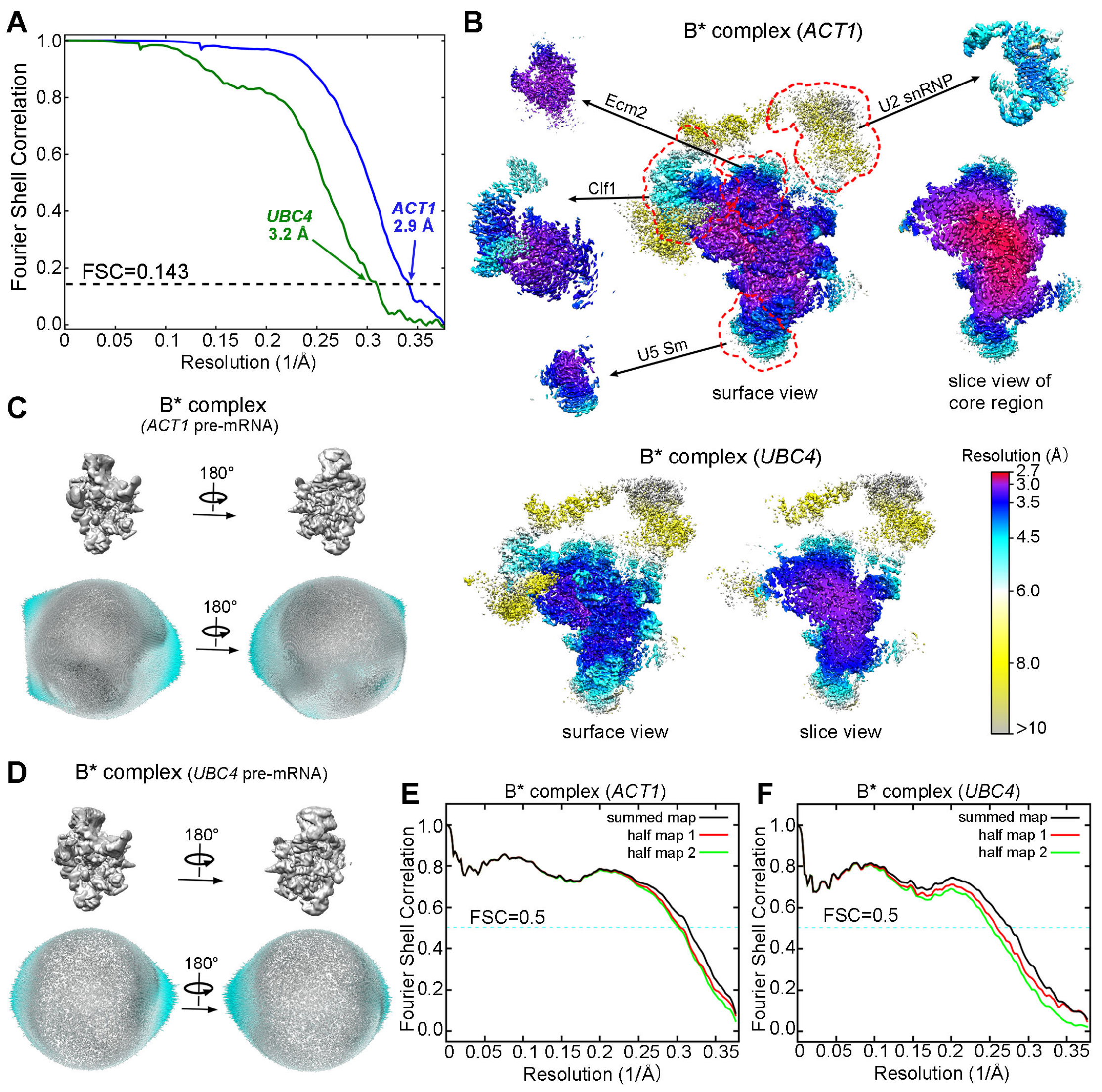


**Figure S4 Cryo-EM analysis of the spliceosomal B$^*$ complexes assembled on the *ACT1* and *UBC4* pre-mRNA from *S. cerevisiae*. Related to Figures 1 & 2** (A) The average resolutions for the *ACT1* B$^*$ complex and the *UBC4* B$^*$ complex are estimated to be 2.9 Å and 3.2 Å, respectively. The resolutions are reported on the basis of the FSC criterion of 0.143. (B) The local resolutions are color-coded for different regions of the *ACT1* B$^*$ complex (upper panel) and the *UBC4* B$^*$ complex (lower panel). The highest resolution of the local EM maps reaches 2.7 Å. (C) Angular distribution of the particles used for reconstruction of the *ACT1* B$^*$ complex. Each cylinder represents one view and the height of the cylinder is proportional to the number of particles for that view. (D) Angular distribution of the particles used for reconstruction of the *UBC4* B$^*$ complex. (E) The FSC curves of the final refined models of the *ACT1* B$^*$ complex versus the individual maps they were refined against (black); of the model refined in the first of the two independent maps used for the gold-standard FSC versus that same map (red); and of the model refined in the first of the two independent maps versus the second independent map (green). The generally similar appearances between the red and green curves indicate that the refinement of the atomic coordinates did not suffer from severe over-fitting. (F) Refinement of the atomic coordinates of the *UBC4* B$^*$ complex did not suffer from severe over-fitting.

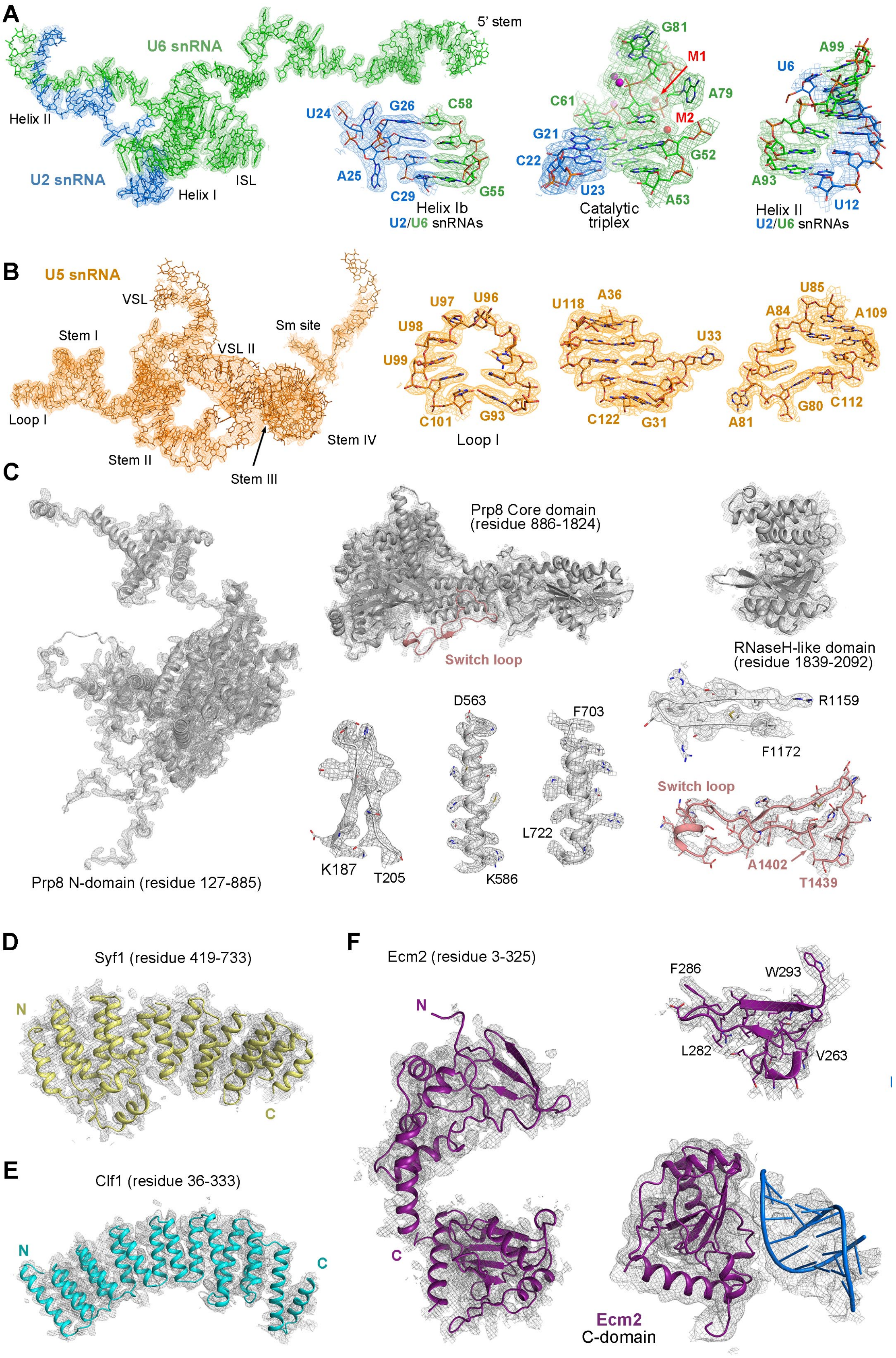


**Figure S5 The EM density maps for the three snRNAs and the four proteins in the B* complex assembled on the *ACT1* pre-mRNA. Related to Figures 1-3** (A)

The EM density maps for U6 snRNA and 30 nucleotides at the 5'-end of U2 snRNA. The overall density map is shown in the left panel. The density maps for helix Ib of the U2/U6 duplex, the catalytic triplex and helix II of U2/U6 duplex are displayed in the middle and right panels, respectively. (B) The EM density maps for U5 snRNA The overall map is shown in the left panel. The density maps for loop I and two representative elements of U5 snRNA are shown in the middle and right panels, respectively. (C) The EM maps for the central component Prp8 are individually shown for the N-domain, the core (which includes the RT Fingers/Palm, Thumb/X, Linker, and endonuclease domain) and the RNaseH-like domain. Close-up views on the EM density maps of some representative secondary structural elements from Prp8 are also shown in the lower right corner. The side chain features for many residues are clearly visible, allowing unambiguous assignment of specific nucleotides and amino acids. (D) The EM density map for residues 419-733 of Syf1. (E) The EM density map for residues 36-333 of Clf1. (F) The EM density maps involving Ecm2. The overall EM density map is shown for residues 3-325 of Ecm2 (left panel). A close-up view is shown for the EM density map of a *de novo* built structural domain (upper right corner). The EM density map is shown for the C-terminal domain of Ecm2 and stem IIb of U2 snRNA (bottom right corner). Unsharpened maps are shown. The Ecm2-U2 interaction has been observed in the C and ILS complexes (Galej et al., 2016; Wan et al., 2016b; Wan et al., 2017).

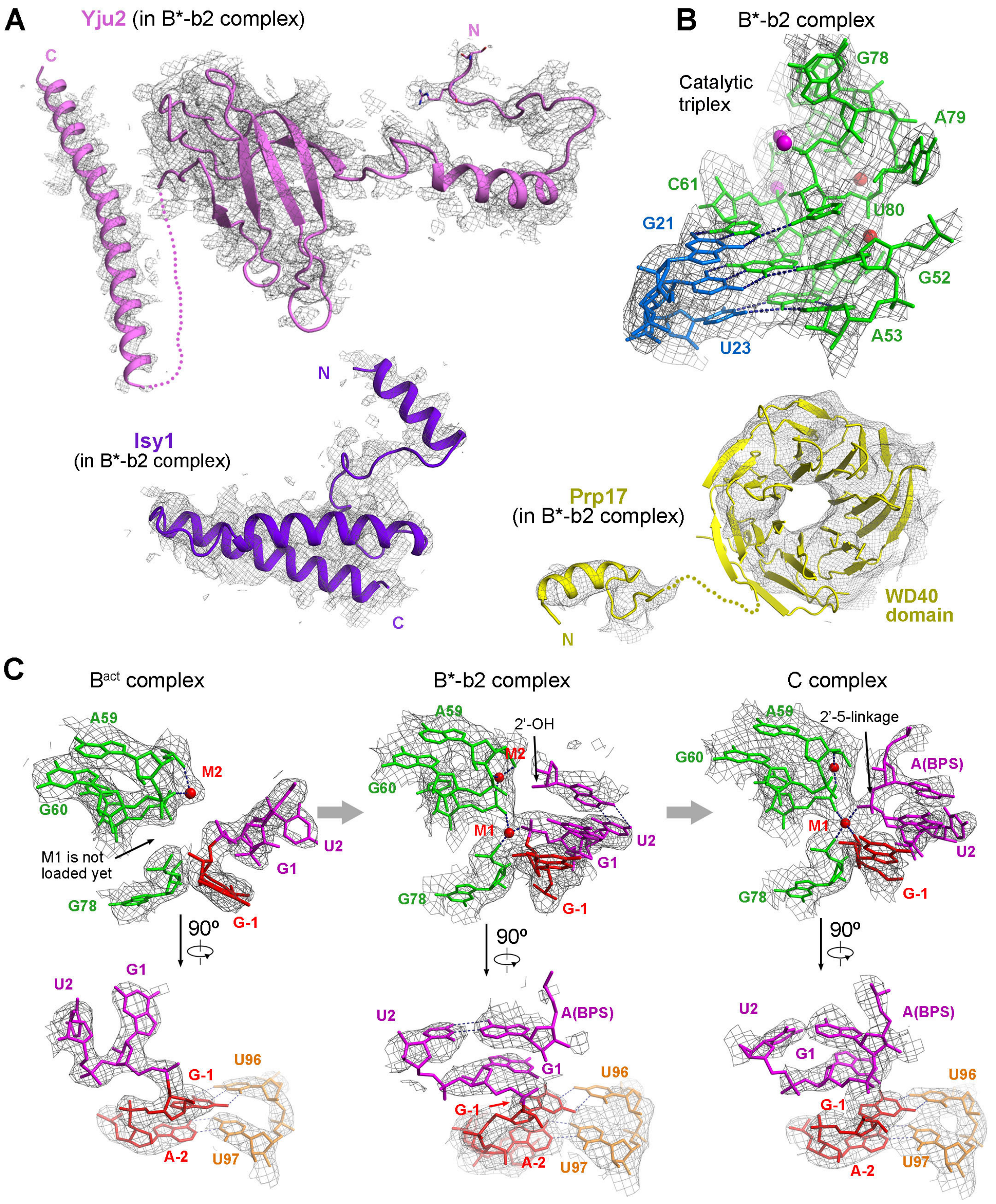


**Figure S6 The EM density maps for the active site components in the B$^{*}$-b2 complex, and the B$^{act}$-to-B$^{*}$-to-C transition. Related to Figure 1, 2, 4 & 5** (A) The EM density maps for Yju2 (left panel), Isy1 (middle panel), and Prp17 (right panel) in the B$^{*}$-b2 complex. The EM density map for Prp17 is low-pass filtered to display the WD40 domain of Prp17. (B) The EM density map of the catalytic triplex in the B$^{*}$-b2 complex. (C) The EM density maps of the active site center in the B$^{act}$

complex (PDB: 5GM6; EMDB: EMD-9524, left panel), the B*-b2 complex (middle panel), and the C complex (PDB: 5GMK; EMDB: EMD-9525, right panel). Two perpendicular views are displayed to highlight the coordination of catalytic metal ions and base-paring interactions with loop I of U5 snRNA.

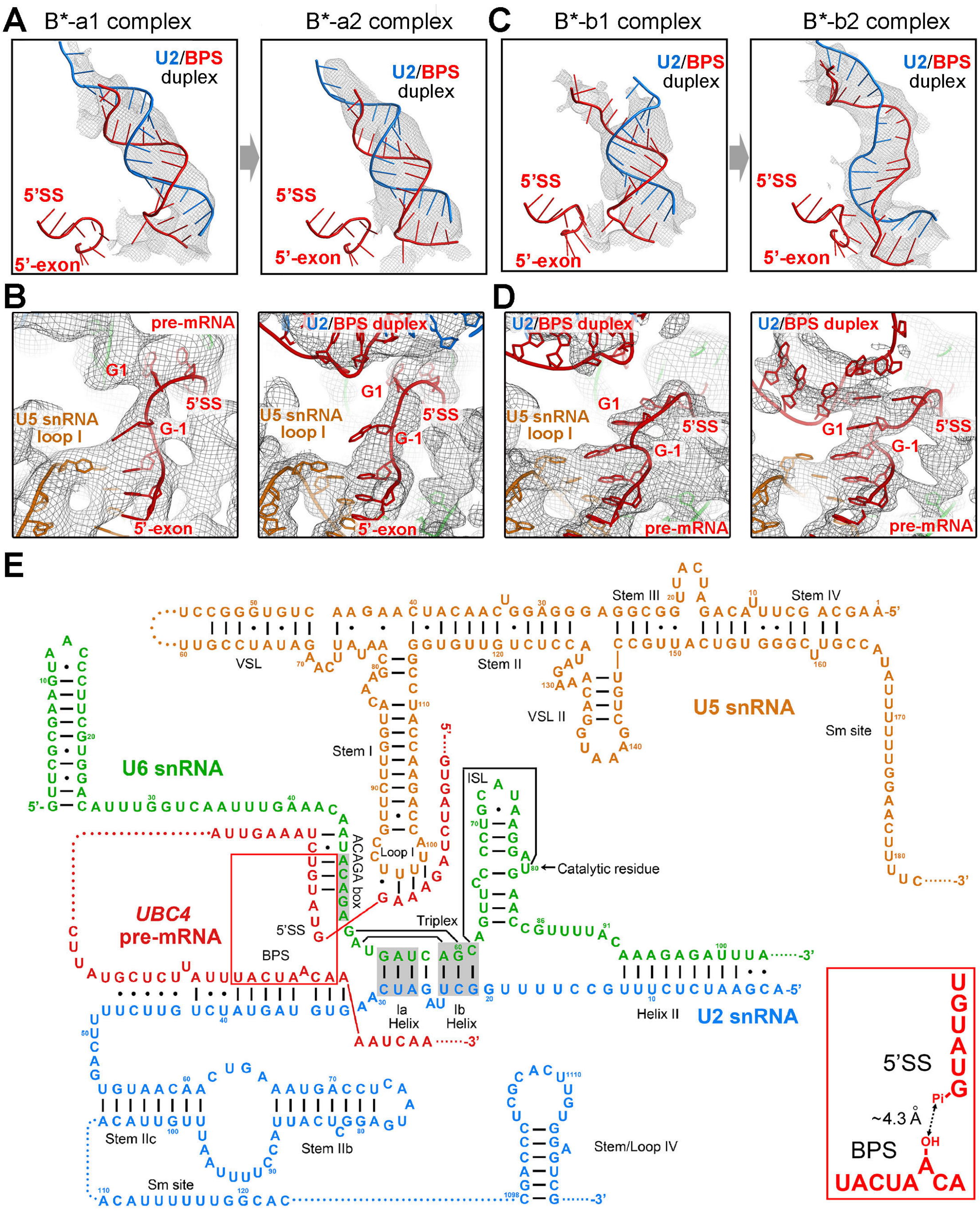


**Figure S7 Secondary structure of the RNA elements in the B*-b2 complex and the EM density maps of the 5'-exon-5'SS region and the U2/BPS duplex in the four B* complexes. Related to Figure 2 & 3** (A) The EM density maps of the U2/BPS duplex for two conformers (a1 and a2) of the *ACT1* B* complex. The duplex is positioned relatively far away from 5'-exon-5'SS in both conformations. (B) The EM density maps of the 5'-exon-5'SS region in the B*-a1 and –a2 complexes. (C)

The EM density maps of the U2/BPS duplex for two conformers (b1 and b2) of the *UBC4* $B^*$ complex. Similar to the two conformations of the *ACT1* $B^*$ complex, the U2/BPS duplex is yet to be docked into the active site in the $B^*$-b1 complex (left panel). With Yju2 loaded, the nucleophile-containing $A_{70}$ of the BPS is already docked into the active site (right panel). (D) The EM density maps of the 5'-exon-5'SS region in the *UBC4* $B^*$-b1 and –b2 complexes. (E) A schematic diagram of the base-pairing interactions among the RNA elements in the $B^*$-b2 complex. Canonical and non-canonical base-pairing interactions are indicated by solid lines and dots, respectively. The flexible sequences yet to be assigned are shown in dotted lines. The RNA elements are colored identically as those in Figure 2A. The inset at the bottom right corner depicts the position of the 5'SS relative to BPS. The distance from the nucleophile (2'-OH of $A_{70}$ in the BPS) to the acceptor (phosphate of $G_1$ at the 5'-end of 5'SS) is approximately 4.3 Å.

### STAR METHODS

#### CONTACT FOR REAGENT AND RESOURCE SHARDING

#### EXPERIMENTAL MODEL AND SUBJECT DETAILS

##### Cell lines

Cef1-CBP tagging was introduced into the *S. cerevisiae* strain 3.2.AID/CRL2101 strain (*MATα, prp2-1, ade2, his3, lys2-801, ura3*, carrying a G360D mutation in Prp2, kindly provided by Dr. Ren-Jang Lin). The resulting yeast cells were cultured in YPD medium to $OD_{600}$ of 2.5~3.5 at 23 °C.

#### METHOD DETAILS

##### CBP tagging in *S. cerevisiae*

Using a PCR-based gene targeting strategy, the carboxyl-terminus of the NTC protein Cef1 was fused to calmodulin binding peptide (CBP) (Figure S1A). The plasmid pF6Aa-CBP-HphMX6 was used as the PCR template for CBP. The PCR products were transformed into the *S. cerevisiae* 3.2.AID/CRL2101 strain (*MATα, prp2-1, ade2, his3, lys2-801, ura3*, carrying a G360D mutation in Prp2, kindly provided by Dr. Ren-Jang Lin) by the lithium acetate method (Gietz and Schiestl, 2007), allowing homologous recombination. Transformants were selected on the YPD (BD) solid medium with hygromycin B (Coolaber Science & Technology). Correct integration of the tag into the genome was confirmed by PCR at the gene level and by Western blots at the protein level. The resulting yeast strain expresses the fusion proteins Cef1-CBP.

**Preparation of the yeast whole cell extract**

72 liters of the Cef1-CBP tagged *S. cerevisiae* 3.2.AID/CRL2101 strain were cultured in the YPD medium at 23 °C to an OD600 of 2.5~3.5. The yeast cells were collected by centrifugation and resuspended in the AGK buffer (10 mM HEPES-KOH, pH 7.9, 200 mM KCl, 1.5 mM $MgCl_2$, 10% (v/v) glycerol and 0.5 mM DTT) in the presence of a protease inhibitor cocktail (final concentrations: 0.5 mM phenylmethylsulphonyl fluoride (PMSF), 2 mM benzamidine, 2.6 μg/ml aprotinin, 1.4 μg/ml pepstatin and 5 μg/ml leupeptin). The cell suspension was dropped into liquid nitrogen to form frozen beads with a diameter of 3-6 mm and pulverized to powder by the SPEX 6870 Freezer Mill. The frozen cell powder was thawed at 4 °C and centrifuged at 18,000g for 30 minutes. The supernatant was centrifuged again at 100,000g for 1 hour, yielding ~100 mL cell extract and a pellet of cell debris. The supernatant was dialyzed twice, each time for 2 hours against 5 liters of Buffer D containing 10 mM HEPES-KOH, pH 7.9, 50 mM KCl, 0.2 mM EDTA, 20% (v/v) glycerol, 0.5 mM DTT and 0.5 mM PMSF, using a 7000 MW cut-off SnakeSkin™ dialysis bag (Thermo Scientific). After dialysis, the whole cell extract was centrifuged at 18,000g for 20 minutes. The supernatant was flash-frozen in 1 mL aliquots and stored at –80 ˚C.

**Expression and purification of Prp2 and Spp2**

The cDNA for the full-length Prp2 or the N-terminally truncated Spp2 (residues 36-185) was amplified from the *Saccharomyces cerevisiae* genomic DNA by PCR and ligated into pET21b vector which contains a C-terminal 8×His tag. The constructs were individually transformed into *Escherichia coli* Rossetta 2 (DE3) strain (Novagen). Cells were grown to an $OD_{600}$ of 0.6~1.0 at 37 °C and induced with 0.4 mM isopropyl-β-D-thiogalactopyranoside (IPTG) to overexpress the target proteins overnight at 18 °C. Cells were harvested by centrifugation, resuspended in the buffer containing 25 mM Tris-HCl, pH 8.0, 500 mM KCl, 1 mM Tris (2-carboxyethyl) phosphine (TCEP) and 5% (v/v) glycerol, and then disrupted by sonication. The recombinant proteins were purified through $Ni^{2+}$-nitrilotriacetate affinity resin (Ni-NTA, Qiagen). The eluted proteins were fractionated through a Heparin column (GE Healthcare) to remove the non-specifically bound nucleic acids. Finally, the proteins were applied to gel filtration (Superdex-200 10/300 GL, GE Healthcare) in the buffer containing 20 mM HEPES-KOH, pH 7.9, 200 mM KCl, 1 mM TCEP and 5% (v/v) glycerol. The peak fractions were analyzed on a 16% SDS PAGE gel (Figure S1B) and stored at -80 °C.

**Preparation of the pre-mRNA**

For the *ACT1* and *UBC4* pre-mRNAs, each was tagged with three tandem MS2-binding RNA aptamers at the 5'-end of the 5'-exon (Figure S1C). The *ACT1* pre-mRNA comprises a 10-nucleotide (nt) 5'-exon, a 309-nt intron and a 178-nt 3'-exon. The *UBC4* pre-mRNA contains a 20-nt 5'-exon, a 95-nt intron and a 20-nt 3'-exon. The 3'-end nucleotides of the 5'-exon in the *ACT1* pre-mRNA are UUCUG, which are non-ideal for base-pairing interactions with the poly-U sequences of U5 loop I. In contrast, the corresponding nucleotides in the *UBC4* pre-mRNA are GAAAG, which should form more stable interactions with U5 loop I. The DNA templates for *in vitro* transcription were generated by PCR, and the RNA transcripts were synthesized by T7 RNA polymerase. The transcriptional products were then resolved on a 6% polyacrylamide denaturing gel and recovered by a diffusion elution buffer containing 200 mM KCl, 50 mM KOAc, pH 6.5, and 2 mM EDTA. Finally, the pre-mRNA was precipitated by ethanol (Sigma) and dissolved in Nuclease-Free Water (Invitrogen).

**Assembly and purification of the yeast $B^{act}$ (△Prp2) complex**

Assembly and purification of the yeast spliceosomal $B^{act}$ (△Prp2) complex was carried out essentially as described (Bao et al., 2017; Warkocki et al., 2009) (Figure S1A). Briefly, the splicing extract was thawed on ice and heated at 35 °C for 30 minutes to inactivate Prp2. The splicing reaction was performed in a volume of 40 mL or multiples thereof and assembled on ice. The reaction contains 60 mM $K\text{-}PO_4$ buffer, pH 7.0, 3% (w/v) PEG8000, 2 mM ATP, 2 mM spermidine, 2.5 mM $MgCl_2$, 10 nM pre-mRNA substrate, 150 nM MS2-MBP and 40% splicing extract. The pre-mRNA substrate was pre-bound to MS2-MBP for 30 minutes on ice. The reaction mixture was incubated for 1 hour at 23 °C and then centrifuged at 18,000g for 15 minutes. The supernatant was incubated with amylose resin (NEB) for 1 hour, allowed to flow through the resin 3 times, and washed using the G150K buffer (10 mM HEPES-KOH, pH 7.9, 150 mM KCl, 1.5 mM $MgCl_2$, 0.01% NP40, 0.5 mM DTT, 0.5 mM PMSF and 5% (v/v) glycerol). The spliceosomal complexes were eluted using 15 mM maltose. This eluted sample was supplemented with 2 mM $CaCl_2$ and then loaded into a calmodulin affinity column (Agilent Technology). The spliceosomal complexes were eluted using the CEB75 buffer (10 mM HEPES-KOH, pH 7.9, 75 mM KCl, 1 mM $Mg(OAc)_2$, 1 mM imidazole, 0.01% NP40 and 2 mM EGTA). The final sample was analyzed on a 6% urea PAGE gel for detection of the RNA elements (Figure S1D).

**Remodeling of the $B^{act}$ (△Prp2) complex by Prp2 and Spp2**

The yeast $B^{act}$ (△Prp2) complex eluted from the calmodulin affinity column (concentrated to ~30 nM) were supplemented with RNasin (0.2 U/μL) and recombinant Prp2 and Spp2 (~900 nM) to a final volume of 1.44 mL in the buffer containing 40 mM K-$PO_4$, pH 7.0, 1.5 mM $MgCl_2$, 0.2 mM EDTA and 5% (v/v) glycerol. Following incubation for 10 minutes on ice, the sample was mixed with 160 μL 10× rescue buffer (200 mM K-$PO_4$, pH 7.0, 14% (v/v) PEG 8000, 10 mM $MgCl_2$, 20 mM ATP) for an hour at 23 °C to allow remodeling (Warkocki et al., 2009). The reaction mixture was supplemented with 2 mM $CaCl_2$ and applied to a calmodulin affinity column. The final sample was analyzed on a 6% urea PAGE gel for detection of the RNA elements (Figure S1D), examined by negative staining EM for particle intactness (Figure S1E), and concentrated for cryo-EM studies (Figure S1F-H).

**EM data acquisition**

Negative staining was carried out essentially as described (Wan et al., 2017). Briefly, the copper grids with a thin layer of carbon film (Zhongjingkeyi Technology) were glow-discharged. A 4-μl aliquot of the sample at ~0.02 mg/ml was applied onto the grid for 1 minute and stored at room temperature. Images were taken on an FEI Tecnai Spirit Bio TWIN microscope operating at 120 kV to verify the sample quality. The Quantifoil R1.2/1.3 grids coated with a thin layer of homemade carbon film were used for cryo-EM specimen preparation. Cryo-EM grids were prepared using Vitrobot Mark IV (FEI Company) at a temperature of 8 °C and a humidity of 100 percent. 4-μl aliquots of the sample at a concentration of ~0.2 mg/mL were applied to glow-discharged grids, blotted for 1.5 seconds, and plunged into liquid ethane cooled by liquid nitrogen. The glow-discharged grids were prepared for 30 seconds using the "Low" setting of the Plasma Cleaner (Harrick, Plasma Cleaner PDC-32G).

The grids were loaded onto an FEI Titan Krios electron microscope equipped with a GIF Quantum energy filter (slit width 20 eV) and operating at 300 kV with a nominal magnification of 105,000×. Images were recorded by a Gatan K2 Summit detector (Gatan Company) using the super-resolution mode, with a pixel size of 0.665 Å. Defocus values were set varied from 1.0 to 2.0 μm. Each image was dose-fractionated to 32 frames with a dose rate of ~8.2 counts/sec/physical-pixel (~6.25 $e^-$

/sec/Å$^2$), total exposure time of 8 seconds, and 0.25 second per frame. AutoEMation 2 (written by Jianlin Lei) was used for all data collection (Lei and Frank, 2005). All 32 frames in each stack were aligned and summed using the whole-image motion correction program MotionCor2 (Zheng et al., 2017) and binned to a pixel size of 1.330 Å. The defocus value of each image was set from 0.8 to 1.8 μm and was determined by Gctf (Zhang, 2016).

**Data Processing**

The EM data processing procedure for the $B^*$ complex assembled on the *ACT1* and *UBC4* pre-mRNA is outlined in Figure S2 and Figure S3, respectively. For the $B^*$ complex assembled on the *ACT1* pre-mRNA, a total of 1,801,049 particles were automatically picked using the deep-learning program DeepPicker (Wang et al., 2016) as previously described (Bai et al., 2018). Particles were extracted and subjected to a guided multi-reference three-dimensional (3D) classification using RELION2.1 (Kimanius et al., 2016; Scheres, 2012). The volumes representing the C complex (EMD-9525), $B^{act}$ complex (EMD-9524), $C^*$ complex (EMD-6684), ILS complex (EMD-6817), and three bad classes were low-pass filtered to 40 Å and used as the initial references (Round 1). To discard particles with other conformation or bad quality, we performed three cycles of global angular searching 3D classification. For each cycle, we merged classes belonging to the $B^*$ complex of the last eleven iterations as the input for the next cycle of global angular searching 3D classification. After that, we performed one cycle of multi-reference local angular searching 3D classification using the references above. 775,243 particles (43.0% of original input) that represent the $B^*$ complex were subjected to a second round (Round 2) of multi-reference local 3D classification with five low-pass filtered references. 2× binned particles (pixel size: 2.66 Å) were used for the classification. Particles from the good classes were combined to yield 555,036 particles (representing 41.7% of the total original particles), which give rise to a 3D reconstruction with an average resolution of 2.9 Å (Figures S2 & S4A). One round of skip-aligned 3D classification with a mask of region around BPS was applied, yielding two major states. Refinements of the two classes of particles produced two reconstructions with overall resolution of 3.6 Å ($B^*$ a1 complex) and 3.2 Å ($B^*$ a2 complex), respectively (Figures S2). Subsequently, refinement with a local mask was individually applied to Ecm2, Clf1, and the U5 Sm ring and U2 snRNP, aiming to improve the local resolutions of the EM maps for the corresponding regions (Figure S4B).

The procedure for data processing of the *UBC4* $B^*$ complex was similar to that for *ACT1* $B^*$ complex. One additional round (Round 3) of local angular searching 3D classification was applied with a soft mask for the core of the $B^*$ complex. We simultaneously performed three parallel 3D classifications and eventually selected 132,125 particles to generate a 3D reconstruction at an overall resolution of 3.2 Å (Figures S3 & S4A; Table S1). Skip-aligned 3D classification with a mask of the region around BPS also yielding two major states. Refinements of them produce two reconstructions with the overall resolution of 3.9 Å ($B^*$-b1 complex) and 3.7 Å (B*-b2 complex (Figures S3).

In the final EM maps of both B$^*$ complexes, the local resolution reaches beyond 3.0 Å in the core region. The resolutions for the regions refined with local masks were further improved (Figure S4B). The angular distributions of the particles used for the final reconstruction of both B$^*$ complexes are reasonable (Figure S4C,D), and the refinement of the atomic coordinates did not suffer from severe over-fitting (Figure S4E, F). The resulting EM density maps display distinguishing features for the amino acid side chains in the core regions of the B$^*$ complex. The RNA elements and their interacting proteins are also well defined by the EM density maps (Figures S5 & S6).

Reported resolutions were calculated on the basis of the FSC 0.143 criterion, with a high-resolution noise substitution method (Chen et al., 2013). Prior to visualization, all density maps were corrected for the modulation transfer function (MTF) of the detector and sharpened by applying a negative B-factor that was estimated using automated procedures (Rosenthal and Henderson, 2003). Local resolution variations were estimated using RELION2.1.

**Model Building and refinement**

Due to a wide range of resolution limits for the various regions of the yeast spliceosomal B$^*$ complexes, we combined homology modeling, rigid docking of known structures, and *de novo* modeling to generate the atomic models (Tables S2 & S3). Identification and docking of the individual components of the B$^*$ complexes were facilitated by the structures of the yeast C (Galej et al., 2016; Wan et al., 2016a), pre-B (Bai et al., 2018) and ILS (Wan et al., 2017) complexes. The protein components that were derived from known structures of the protein data bank (PDB) are summarized in Tables S2 & S3. These structures were docked into the density map using COOT (Emsley and Cowtan, 2004) and fitted into density using CHIMERA (Pettersen et al., 2004).

The atomic coordinates of U6 snRNA, U2 snRNP core, U5 snRNP, and the NTC and NTR complexes from the yeast C complex (Wan et al., 2016a) were docked into the density maps of the yeast B* complexes and were manually adjusted using COOT (Emsley and Cowtan, 2004). The pre-mRNA was manually adjusted and extended using COOT. The U1 Sm ring from the yeast pre-B complex was used as the initial model for U2 and U5 Sm ring. The VSL II, Stem III and Stem IV of U5 snRNA were built using those in the pre-B complex structure as template (Bai et al., 2018). The atomic coordinates of Ecm2, Syf1 and Clf1 from the yeast C complex (Wan et al., 2016a) were docked into to the B* complex and manually adjusted and extended. The final atomic models of both the B* complexes were refined against their respective EM maps using PHENIX in real space (Adams et al., 2010) and secondary structure restraints that were generated by ProSMART (Nicholls et al., 2014). Overfitting of the overall model was monitored by refining the model in one of the two independent maps from the gold-standard refinement approach, and testing the refined model against the other map (Amunts et al., 2014). The structures of the yeast B* complexes were validated through examination of the Molprobity scores and statistics of the Ramachandran plots (Table S1). Molprobity scores were calculated as described (Davis et al., 2007) .

#### QUANTIFICATION AND STATISTICAL ANALYSIS

Resolution estimations of cryo-EM density maps are based on the 0.143 Fourier Shell Correlation (FSC) criterion (Chen et al., 2013; Rosenthal and Henderson, 2003).

#### DATA AND SOFTWARE AVAILABILITY

##### Data Resources

Atomic coordinates and EM density maps of the *S. cerevisiae ACT1* $B^*$ complex (PDB: xxxx; EMDB: EMD-xxxx) and *UBC4* $B^*$ complex (PDB: xxxx; EMDB: EMD-xxxx) have been deposited with the Protein Data Bank (http://www.rcsb.org) and the Electron Microscopy Data Bank (https://www.ebi.ac.uk/pdbe/emdb/).

**Table S1 Cryo-EM data collection and refinement statistics of the B* complex from *Saccharomyces cerevisiae* (*S. cerevisiae*).**

| | B* complex (*ACT1* pre-mRNA) | B* complex (*UBC4* pre-mRNA) |
|---|---|---|
| **Data collection** | | |
| EM equipment | FEI Titan Krios | FEI Titan Krios |
| Voltage (kV) | 300 | 300 |
| Detector | Gatan K2 | Gatan K2 |
| Pixel size (Å) | 1.330 | 1.330 |
| Electron dose (e-/Å$^2$) | 49.3 | 49.3 |
| Defocus range (μm) | 1.0~2.0 | 1.0~2.0 |
| **Reconstruction** | | |
| Software | RELION 2.1 | RELION 2.1 |
| Number of used Particles | 555,036 | 132,125 |
| Accuracy of rotation (°) | 0.532 | 0.563 |
| Accuracy of translation (pixels) | 0.300 | 0.327 |
| Final Resolution (Å) | 2.9 | 3.2 |
| Map sharpening B-factor (Å$^2$) | -89.8036 | -98.7873 |
| **Model building** | | |
| Software | Coot | Coot |
| **Model composition** | | |
| Protein residues | 8982 | 9168 |
| RNA nucleotides | 549 | 548 |
| GTP | 1 | 1 |
| IP6 | 1 | 1 |
| **Refinement** | | |
| Software | PHENIX1.13 | PHENIX1.13 |
| R-factor | 0.3249 | 0.3280 |
| **Validation** | | |
| R.m.s deviations | | |
| Bonds length (Å) | 0.007 | 0.010 |
| Bonds Angle (°) | 1.060 | 1.142 |
| Ramachandran plot statistics (%) | | |
| Preferred | 94.12 | 93.25 |
| Allowed | 5.47 | 6.29 |
| Outlier | 0.41 | 0.46 |
| RNA validation | | |
| Correct sugar pucker (%) | 95.72 | 97.65 |
| Good Backbone conformations (%) | 62.65 | 69.02 |
| Molprobity score | 2.14 | 2.10 |

#### Table S2 Summary of model building for the B* complex (*ACT1* pre-mRNA).

| | Components *S. cerevisiae*/*S. pombe*/Human | Total length | Modeled domain/region | Modeling template | Modeling | Resolution (Å) | Chain ID |
|---|---|---|---|---|---|---|---|
| **Pre-mRNA** | *ACT1* pre-mRNA | 497 | 5'exon, (-12)-(-1)<br>intron (1-18, 247-276) | - | *de novo* | 2.7~6 | B |
| **U5 snRNP** | Prp8/Spp42/Prp8 | 2413 | N-domain (127-885)<br>RT finger/palm (886-1253)<br>Thumb/X (1254-1377)<br>Linker (1378-1650)<br>Endonuclease (1651-1829)<br>RNaseH-like (1840-2085) | 5GMK | Docked & rebuilt | 2.7~4.5 | A |
| | U5 snRNA | 214 | 1-183 | 5GMK/ 5ZWM | Docked & rebuilt | 2.7~4.5 | D |
| | Snu114/Cwf10/Snu114 | 1008 | 67-985<br>GTP | 5GMK | Docked & rebuilt | 2.7~3.5 | C |
| | SmB,D1,D2,D3,E,F,G | | Sm fold | 5ZWN | Rigid docking | 3.0~10.0 | g-m |
| **U6 snRNA** | U6 snRNA | 112 nt | 1-103 | 5GMK | Docked & rebuilt | 2.7~4.5 | E |
| **U2 snRNP** | U2 snRNA | 1175 nt | 1-52<br>54-123<br>1082-1169 | 5GMK/ 5LJ3 | Docked & rebuilt<br>Rigid docking<br>Rigid docking | 3.0~8.0<br>10.0~20.0<br>10.0~20.0 | L |
| | Msl1/Msl1/U2-B" | 111 | RRM domain | 5GMK | Rigid docking | 8.0~20.0 | a |
| | Lea1/Lea1/U2-A' | 238 | LRR domain | 5GMK/ 5LJ3 | Docked & adjusted | 8.0~20.0 | b |
| | SmB,D1,D2,D3,E,F,G | | Sm fold | 5ZWN | Rigid docking | 8.0~20.0 | e, s, u, w-z |
| **NTC** | Prp19 | 503 | 1-139 | 5GMK | Rigid docking | ~20 | o-r |
| | Cef1/Cdc5/Cdc5 | 590 | Myb Domain (9-111)<br>145-253<br>332-587 | | Docked & rebuilt<br>Docked & rebuilt<br>Rigid docking | 3.0~4.5<br>3.0~4.5<br>~20 | c |
| | Snt309/Cwf7/Spf27 | 175 | 1-175 | | Rigid docking | ~20 | t |
| | Syf1/Cwf3/Syf1 | 859 | 47-418<br>419-734<br>734- | | Idealized α helix<br>Docked & rebuilt<br>Idealized α helix | 3.0~15.0 | v |
| | Clf1/Cwf4/Syf3 | 687 | 36-333<br>334-641 | | Docked & rebuilt<br>Idealized α helix | 3.0~4.0<br>5.0~15.0 | d |
| | Syf2 | 215 | 92-211 | | Docked & rebuilt | 3.5~4.5 | I |
| | Isy1/Cwf12/Isy1 | 235 | 27-96 | | Docked & rebuilt | 4.5~8.0 | H |
| **NTC Related proteins** | Prp46/Prp5/PRL1 | 451 | 95-128, 429-451<br>WD40 domain (120-428) | 5GMK | Docked & rebuilt<br>Docked & rebuilt | 3.0~4.5 | O |
| | Prp45/Prp45/SKIP | 379 | 28-247 | | Docked & rebuilt | 3.0~4.5 | P |
| | Ecm2/Cwf5/RBM22 | 364 | 3-125, 202-227, 239-264<br>126-146, 179-201, 265-325 | | Docked & rebuilt<br>*de novo* | 3.0~4.5 | Q |
| | Cwc2/Cwf2/RBM22 | 339 | 1-261 | | Docked & rebuilt | 3.0~4.5 | R |
| | Cwc15/Cwf15/Ad002 | 175 | 4-41, 126-175 | | Docked & rebuilt | 3.0~4.5 | S |
| | Bud31/Cwf14/G10 | 157 | 1-157 | | Docked & rebuilt | 3.0~4.5 | T |
| **Splicing Factors** | Cwc21/Cwf21/SRRM2 | 135 | 2:28 | 5GMK | Docked & rebuilt | 3.5~4.5 | J |
| | Cwc22/Cwf22/Cwc22 | 577 | MIF4G domain (8:247)<br>MA3 domain (279:485) | -<br>5GMK | Rigid docking<br>Docked & rebuilt | 3.5~10<br>3.5~4.5 | Z |
| | Prp17/Prp17/CDC40 | 455 | 51:73<br>WD40 domain (151:454) | 5GMK<br>5XJC | Docked & rebuilt<br>Rigid docking | 3.0~4.5<br>3.5~8.0 | n |

Under the column labeled "Components", proteins from *S. cerevisiae*, *S. pombe*, and human are colored black, red, and green, respectively. If the proteins from all three species have the same name, only a single name in black is indicated.

#### Table S3 Summary of model building for the B* complex (*UBC4* pre-mRNA).

| | Components *S. cerevisiae*/*S. pombe*/Human | Total length | Modeled domain/region | Modeling template | Modeling | Resolution (Å) | Chain ID |
|---|---|---|---|---|---|---|---|
| **Pre-mRNA** | *UBC4* pre-mRNA | 497 | 5'exon, (-12)-(-1)<br>intron (1-16, 51-79) | - | *de novo* | 3.0~6 | B |
| **U5 snRNP** | Prp8/Spp42/Prp8 | 2413 | N-domain (127-885)<br>RT finger/palm (886-1253)<br>Thumb/X (1254-1377)<br>Linker (1378-1650)<br>Endonuclease (1651-1829)<br>RNaseH-like (1840-2085) | 5GMK | Docked & rebuilt | 3.0~4.5 | A |
| | U5 snRNA | 214 | 1-183 | 5GMK/<br>5ZWM | Docked & rebuilt | 3.0~4.5 | D |
| | Snu114/Cwf10/Snu114 | 1008 | 67-985<br>GTP | 5GMK | Docked & rebuilt | 2.7~3.5 | C |
| | SmB,D1,D2,D3,E,F,G | | Sm fold | 5ZWN | Rigid docking | 3.5~10.0 | g-m |
| **U6 snRNA** | U6 snRNA | 112 nt | 1-103 | 5GMK | Docked & rebuilt | 3.0~4.5 | E |
| **U2 snRNP** | U2 snRNA | 1175 nt | 1-52<br>54-123<br>1082-1169 | 5GMK/<br>5LJ3 | Docked & rebuilt<br>Rigid docking<br>Rigid docking | 3.0~8.0<br>10.0~20.0<br>10.0~20.0 | L |
| | Msl1/Msl1/U2-B" | 111 | RRM domain | 5GMK | Rigid docking | 8.0~20.0 | a |
| | Lea1/Lea1/U2-A' | 238 | LRR domain | 5GMK/<br>5LJ3 | Docked & adjusted | 8.0~20.0 | b |
| | SmB,D1,D2,D3,E,F,G | | Sm fold | 5ZWN | Rigid docking | 8.0~20.0 | e, s, u, w-z |
| **NTC** | Prp19 | 503 | 1-139 | 5GMK | Rigid docking | ~20 | o-r |
| | Cef1/Cdc5/Cdc5 | 590 | Myb Domain (9-111)<br>145-253<br>332-587 | | Docked & rebuilt<br>Docked & rebuilt<br>Rigid docking | 3.5~4.5<br>3.5~4.5<br>~20 | c |
| | Snt309/Cwf7/Spf27 | 175 | 1-175 | | Rigid docking | ~20 | t |
| | Syf1/Cwf3/Syf1 | 859 | 47-418<br>419-734<br>734- | | Idealized α helix<br>Docked & rebuilt<br>Idealized α helix | 3.5~15.0 | v |
| | Clf1/Cwf4/Syf3 | 687 | 36-333<br>334-641 | | Docked & rebuilt<br>Idealized α helix | 3.5~4.0<br>5.0~15.0 | d |
| | Syf2 | 215 | 92-211 | | Docked & rebuilt | 3.5~4.5 | I |
| | Isy1/Cwf12/Isy1 | 235 | 2-96 | | Docked & rebuilt | 4.5~8.0 | H |
| **NTC Related proteins** | Prp46/Prp5/PRL1 | 451 | 95-128, 429-451<br>WD40 domain (120-428) | 5GMK | Docked & rebuilt<br>Docked & rebuilt | 3.5~4.5 | O |
| | Prp45/Prp45/SKIP | 379 | 28-247 | | Docked & rebuilt | 3.5~4.5 | P |
| | Ecm2/Cwf5/RBM22 | 364 | 3-125, 202-227, 239-264<br>126-146, 179-201, 265-325 | | Docked & rebuilt<br>*de novo* | 3.5~4.5 | Q |
| | Cwc2/Cwf2/RBM22 | 339 | 1-261 | | Docked & rebuilt | 3.5~4.5 | R |
| | Cwc15/Cwf15/Ad002 | 175 | 4-41, 126-175 | | Docked & rebuilt | 3.5~4.5 | S |
| | Bud31/Cwf14/G10 | 157 | 1-157 | | Docked & rebuilt | 3.5~4.5 | T |
| **Splicing Factors** | Cwc21/Cwf21/SRRM2 | 135 | 2:28 | 5GMK | Docked & rebuilt | 3.5~4.5 | J |
| | Cwc22/Cwf22/Cwc22 | 577 | MIF4G domain (8:247)<br>MA3 domain (279:485) | -<br>5GMK | Rigid docking<br>Docked & rebuilt | 3.5~10<br>3.5~4.5 | Z |
| | Prp17/Prp17/CDC40 | 455 | 51:73<br>WD40 domain (151:454) | 5GMK<br>5XJC | Docked & rebuilt<br>Rigid docking | 3.5~4.5<br>3.5~8.0 | n |
| | Yju2/ Cwf16/Yju2 | 278 | 2-37<br>38-116<br>166-211 | 5GMK<br>5GMK<br>5Y88 | Docked & rebuilt<br>Rigid docking<br>Docked & adjusted | 3.5~4.5<br>4.5~8.0<br>4.5~8.0 | F |

Under the column labeled "Components", proteins from *S. cerevisiae*, *S. pombe*, and human are colored black, red, and green, respectively. If the proteins from all three species have the same name, only a single name in black is indicated.
